# Engineering a Single Extracellular Vesicle Protein and RNA Assay (^siEV^PRA) via In Situ Fluorescence Microscopy in a UV Micropatterned Array

**DOI:** 10.1101/2022.08.05.502995

**Authors:** Jingjing Zhang, Xinyu Wang, Xilal Y. Rima, Luong T. H. Nguyen, Kristin Huntoon, Yifan Ma, Nicole Walters, Kwang Joo Kwak, Min Jin Yoon, Daeyong Lee, Yifan Wang, Jonghoon Ha, Kelsey Scherler, Shannon Fallen, Inyoul Lee, Andre F. Palmer, Wen Jiang, Kai Wang, Betty Y.S. Kim, L. James Lee, Eduardo Reátegui

## Abstract

The physical and molecular heterogeneity of extracellular vesicles (EVs) confounds bulk biomarker characterization, thus encouraging the development of novel assays capable of profiling EVs at a single-vesicle resolution. Here, we present a single EV (siEV) protein and RNA assay (^siEV^PRA) to simultaneously detect proteins, messenger RNAs (mRNAs), and microRNAs (miRNAs) in siEVs. The ^siEV^PRA consists of an array of microdomains embedded on a polyethylene glycol (PEG)-coated glass surface produced via UV photopatterning, functionalized with antibodies to target siEV subpopulations. Fluorescently labeled antibodies and RNA-targeting molecular beacons (MBs) were used to generate signals for proteins, mRNAs, and miRNAs on siEVs detected by total internal reflection fluorescence microscopy (TIRFM), outperforming the sensitivities of ELISA and PCR by three orders of magnitude. Using the ^siEV^PRA, we analyzed EVs harvested from glioblastoma (GBM) cell lines and demonstrated vesicular heterogeneity in protein, mRNA, and miRNA expression through colocalization analyses, and validated the results by bulk RNA sequencing. We further demonstrated the clinical utility of the ^siEV^PRA by detecting different mRNAs and miRNAs associated with GBM in patient samples. Together, these results indicate that the ^siEV^PRA provides an effective platform to investigate the heterogeneity of proteins and RNAs in subpopulations of EVs.

## 1. Introduction

Extracellular vesicles (EVs) are small membranous vesicles released by cells and are present in bodily fluids^1^. EVs have been shown to play a role in different biological processes that span from physiological tissue regulation to pathogenic injury and organ remodeling^2^. Despite the potential use of EVs in the clinic as diagnostic and therapeutic tools for different diseases, current methods for isolating and characterizing EVs are technically challenging^3,4^. Isolation methods are usually cumbersome and irreproducible, while characterization relies on techniques including western blotting (WB), enzyme-linked immunosorbent assay (ELISA), polymerase chain reaction (PCR), next-generation sequencing (NGS), and mass spectroscopy (MS) which provide an average measurement of the nucleic acid and protein content. Consequently, with these characterization techniques, EVs are physically broken down to obtain their internal contents, whereby essential molecular information of tissue-specific single EVs (siEVs) can be lost. EVs are highly heterogeneous in molecular composition, with their proteins, RNAs, DNAs, lipids, and metabolites reflecting their tissue of origin^5,6^. Investigating the molecular information within siEVs is necessary to understand the effects of EV-membrane proteins and vesicular cargo on EV-mediated intercellular signaling in diseases such as cancer. EVs have been shown to promote drug resistance^7^, immunosuppression^8^, the epithelial-to-mesenchymal transition (EMT)^9^, and metastasis^10^. Therefore, there is a critical need to develop technologies that provide an accurate and efficient analysis of the molecular content within siEVs.

Several analytical methods have been reported to quantify the physical and molecular characteristics of siEVs. Nanoparticle tracking analysis (NTA) and tunable resistive pulse sensing (TRPS) are routinely used to measure the size and concentration of siEVs, with the minimum detectable size of EVs in the 70 −100 nm range^11^. However, NTA and TRPS lack specificity to characterize tissue-specific siEVs. Flow cytometry can detect siEVs as small as 40 nm, incorporating fluorescent protein detection^12^. However, reduced multiplexed capability, aggregation or swarmingof EVs due to the required concentrations, and extensive calibration requirements have limited their use. On the other hand, surface and cargo proteins have been characterized in siEVs using nano-plasmonic and interferometric biosensors^13–15^. Moreover, antibody-DNA conjugates incorporating a random tag sequence in a proximity barcoding assay with NGS have been used to profile different proteins simultaneously in siEVs^16^. Although these promising technologies have demonstrated their ability to resolve subpopulation of siEVs from different tissues, the complex cargo of EVs, such as nucleic acids, still requires strategies that enable different types of molecular cargo quantification.

Recently, super-resolution microscopy methods have been used to detect and quantify single proteins and nucleic acids at the sub-vesicular level to unravel the heterogeneity of EVs derived from biofluids^17,18^. Quantitative single-molecule localization microscopy (qSMLM) can characterize the size and membrane protein content of siEVs from plasma^19^. Stochastic optical reconstruction microscopy (STORM) combined with total internal fluorescence microscopy (TIRFM) has improved the signal-to-noise ratio and reduced the imaging time of siEVs^18,20^. However, the nucleic acid cargo analysis of siEVs involves intricate chemistries that usually alter the native structure of EVs, producing high background signal levels, thus limiting the use of super-resolution microscopy to analyze highly expressed RNA biomarkers in siEVs^17^. Moreover, the low-throughput nature of these techniques has also limited their broad dissemination for clinical use^18^.

Here, we describe the single EV protein and RNA assay (^siEV^PRA), capable of multiplexing protein and RNA biomarker detection at a single-vesicle resolution. The assay consists of an array of microdomains patterned on a polyethylene glycol (PEG)-coated glass surface using UV light with a digital-micromirror device (DMD) that allows maskless photopatterning. The arrayed surface is functionalized with antibodies against EV-specific epitopes, such as tetraspanins, ADP-ribosylation factor 6 (ARF6), and Annexin A1 to immobilize subpopulations of siEVs onto distinct positions. Fluorescently labeled antibodies and RNA-targeting molecular beacons (MBs) are used to generate signals for proteins, mRNAs, and miRNAs on siEVs detected by TIRFM and quantified via automatic image acquisition. The ^siEV^PRA exceeds the detection limit for both ELISA and PCR by three orders of magnitude without tedious EV lysis extraction procedures. The ability of the ^siEV^PRA to multiplex various biomarkers within and across biomolecule species enables complex EV heterogeneity analyses such as simultaneous protein and RNA detection of up to 9 different biomarkers in siEVs (4 proteins and 5 RNAs) enriched with different capture antibodies. In this work, the ^siEV^PRA was extended to investigate subpopulations of EVs from glioblastoma (GBM) cell lines to determine the heterogeneity of different RNAs, confirmed with bulk RNA sequencing. Next, we established the clinical utility of the ^siEV^PRA by validating the expression of different mRNAs and miRNAs associated with GBM in siEVs. The siEV analysis of serum from GBM patients demonstrated that distinctive RNA signals were obtained when compared to healthy controls. To the best of our knowledge, this is the first assay that enables the simultaneous and low-dose profiling of protein, miRNA, and mRNA on siEVs, lending unique applications for liquid biopsies and therapeutics.

## 2. Results and Discussion

### 2.1. Single-EV analysis with the ^siEV^PRA

Our device was fabricated with the PRIMO optical module (**Fig. 1A**). A glass coverslip was coated with poly-L-lysine (PLL) through physisorption and methoxy-poly(ethylene glycol)-succinimidyl valerate (mPEG-SVA) was covalently bound to the surface through N-hydroxysuccinimide (NHS) chemistry creating a non-biofouling coating. A five-by-five array of 20-μm diameter circles was cleaved from the mPEG monolayer via UV projections translated by a DMD in the presence of 4-benzoylbenzyl-trimethylammonium chloride (PLPP) as a photoactivator. The level of photoscission correlates to both the grayscale value of a digital template and the UV dose^21^. A 50 % grayscale value and a 20 mJ/mm^2^ dose were selected as they rendered the highest relative fluorescent intensity (RFI) to capture siEVs relative to the control (phosphate buffered saline, PBS) and minimized non-specific binding within the microdomains (**Supplementary Table 1**). The optimized grayscale value and dose demonstrated homogenous adsorption of NeutrAvidin (NA) with specificity to the photocleaved surfaces (**Supplementary Fig. 1**). Given the uniformity of the NA layer, biotinylated antibodies against CD63 and CD9, epidermal growth factor receptor (EGFR), ARF6, and Annexin A1, which are present as membrane proteins on EVs, were patterned in the microdomains to tether siEVs selectively (**Fig. 1A**).

**Figure 1:**
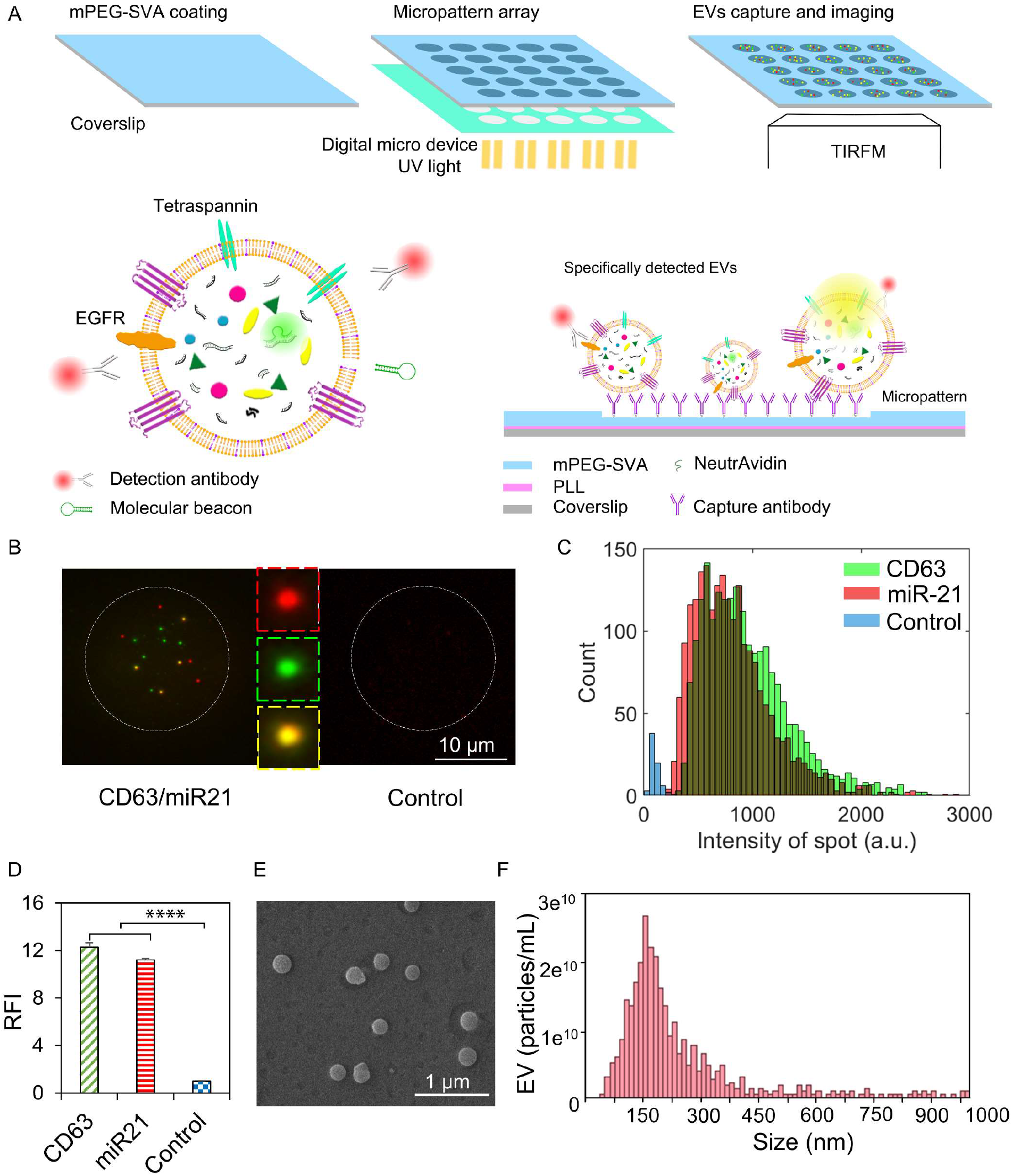
Single-EV detection with the ^siEV^PRA integrated assay. **(A)** Schematicm representation of the device fabrication with the PRIMO optical module to photoetch microdomains via DMD-based UV projections, where NeutraAvidin (NA) was physisorbed to tether biotinylated antibodies against epitopes on the surfaces of siEV. **(B)** CD63 (green dots) and miR-21 (red dots) on Gli36-derived siEVs are detected and colocalized (yellow dots) with the ^siEV^PRA. The control sample (no EVs) demonstrates a negligible fluorescent signal. **(C)** Images are quantified as statistical distributions to depict the expression of CD63 and miR-21 at a single-vesicle level for the different samples. **(D)** Quantifying relative fluorescence intensity (RFI) for CD63, miR-21 in siEVs captured in the device. The control sample demonstrates different counts (n = 3, error bars indicate the standard deviation). **(E)** SEM of a typical sample confirms the presence of siEVs tethered to the surface of the device. **(F)** Gli36-derived siEV size distribution and concentration measured by TRPS demonstrate heterogeneity in particle size.

To test the presence of siEVs on the microdomains, a fluorescently labeled antibody against CD63 and an MB targeting miR-21, an abundant EV-enveloped miRNA, was used as detection probes and visualized via TIRFM. Each green fluorescent spot represented a siEV expressing CD63, and each red fluorescent spot represented a siEV carrying miR-21. Each yellow spot demonstrated the colocalization of both biomarkers at a single-vesicle resolution. Conversely, fluorescent signals in control samples were significantly lower, indicating the ability of the ^siEV^PRA to multiplex different biomolecule species in siEVs (**Fig. 1B**). Furthermore, acquired TIRFM images could be quantified as statistical fluorescent signal distributions to analyze biomarker expression on siEVs (**Fig. 1C**) or to quantify the RFI of the sample (**Fig. 1D**), which revealed an RFI of 12.16 ± 0.50 and 11.26 ± 0.08 for siEVs relative to the control samples (ANOVA, *p* < 0.0001 for CD63 and miR-21). After EV immobilization, scanning electron microscopy (SEM) was performed on the device to further validate the fluorescent signals observed on the microdomains as originating from siEVs. The SEM images revealed single, round vesicles, confirming the presence of siEVs tethered on the microdomains (**Fig. 1E**). TRPS measurements on the EV samples used for the ^siEV^PRA demonstrated a mean-siEV diameter of 150 nm, consistent with the size of the vesicles observed by SEM (**Fig. 1F**). Thus, the ^siEV^PRA successfully captures siEVs in distinct surface array positions and multiplexes protein and RNA signals via immunoaffinity and MB RNA hybridization, respectively.

### 2.2. Specificity and sensitivity of RNA and protein detection in siEVs

Although there are methods available to detect proteins on siEVs, detecting RNA without altering or damaging the integrity of the vesicles remains a challenge^22^. To detect membrane-enveloped RNAs, MBs were diluted in a tris-EDTA (TE) buffer to partially permeate the lipid bilayer of the EVs, allowing the MBs to penetrate the membrane and hybridize with the desired RNA sequences^23^. The changes in EV concentration when incubating in the TE buffer and PBS were negligible (ANOVA, *p* = 0.65), implying the extent of permeabilization by the TE buffer was not detrimental to the integrity of the EVs (**Supplementary Fig. 2**). To ensure the specificity of the MBs to the desired RNA targets on the ^siEV^PRA, miR-21, a human miRNA, and miR-39, a non-human miRNA abundant in *Caenorhabditis elegans*^24^, were tested in siEVs derived from Gli36 cells, a human glioma cell line. Gli36-derived EVs detected with MBs targeting miR-21 exhibited single fluorescent spots within the microdomain when diluted in the TE buffer (**Supplementary Fig. 3A**). The MB formulation diluted in the TE buffer produced a fluorescent signal 6.30 ± 0.50 times higher than the formulation diluted in PBS when applied to the immobilized siEVs (ANOVA, *p* < 0.0001), indicating the necessity for partial permeabilization. Furthermore, the siEV signals obtained from partial permeabilization were 9.65 ± 1.28 times higher than MBs diluted solely in the TE buffer (ANOVA, *p* = 0.0054) and 9.80 ± 1.30 times higher than MBs diluited solely in PBS (ANOVA, *p* < 0.0001), ensuring the specificity of the MBs to detect RNAs in siEVs. In contrast, Gli36-derived EVs detected with MBs targeting non-human miR-39 within the TE buffer and PBS demonstrated a negligible difference when compared to their respective controls (ANOVA, *p* = 0.62 for TE and *p* = 0.68 for PBS), thus proving the ability of the ^siEV^PRA to target specific RNA sequences (**Supplementary Fig. 3B**).

To evaluate the robustness of RNA specificity using the ^siEV^PRA, Gli36 cells were transfected via electroporation with cel-miR-54, cel-miR-39, and cel-miR-238 plasmids, which are non-human miRNAs (**Supplementary Fig. 4A**)^24–26^. EVs harvested from the transfected cells were then detected with MBs targeting miR-39, miR-54, and miR-238. The engineered siEVs loaded with non-human miRNAs were successfully detected as single fluorescent spots within the microdomains when the MBs targeted the corresponding miRNA. In contrast, control samples showed a negligible number of fluorescent spots (**Fig. 2A**). To ascertain a lack of cross-reactivity between the MBs and the other non-human miRNA, the three different engineered EVs were tested against all the MBs targeting the non-human miRNA. Only the MBs targeting the corresponding nonhuman miRNA loaded within the engineered EVs could be detected, whereas all disparate MBs presented a background level of fluorescent spots (**Supplementary Fig. 4B**). Similarly, EVs collected from healthy donor serum presented few fluorescent spots (**Supplementary Fig. 4B**). Specifically, miR-54-enriched EVs detected by MBs targeting miR-54 produced a fluorescent signal 9.43 ± 1.68 times higher than the average of the controls (ANOVA, p < 0.0001), miR-39-enriched EVs detected by MBs targeting miR-39 produced a fluorescent signal 9.10 ± 2.07 times higher than the average of the controls (ANOVA, p < 0.0001), and miR-238-enriched EVs detected by MBs targeting miR-238 produced a fluorescent signal 8.73 ± 2.52 times higher than the average of the controls (ANOVA, p < 0.0001) (**Fig. 2B**). Furthermore, the ^siEV^PRA was capable of discriminating between the EVs transfected with varying plasmid concentrations, demonstrating the sensitivity of the assay to quantify nucleic acid concentrations within siEVs (**Fig. 2C**).

**Figure 2:**
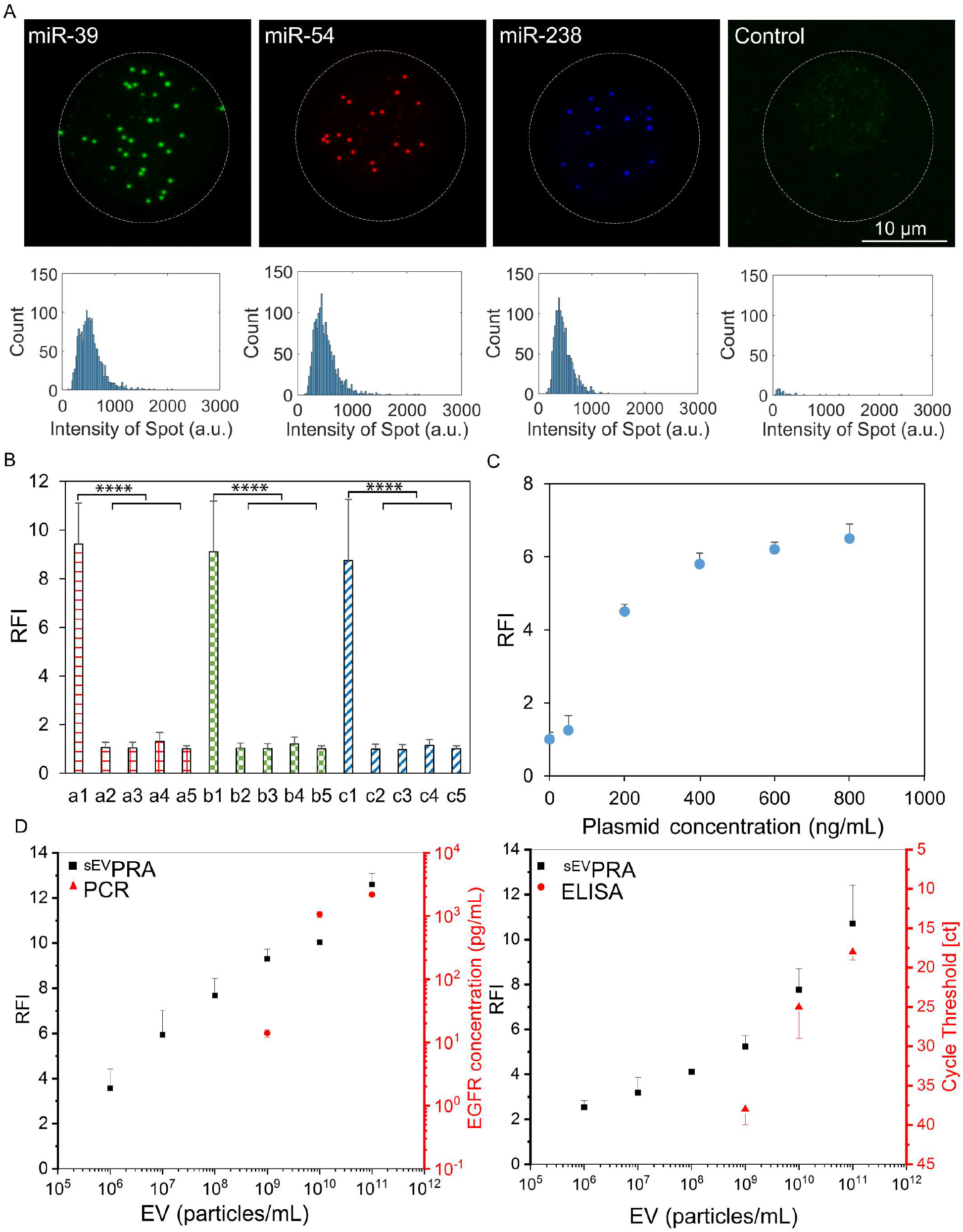
Specificity and sensitivity of RNA and protein detection. **(A)** siEVs loaded with miR-54, miR-39, and miR-238 are detected by the corresponding MBs targeting miR-39 (green), miR-54 (red), and miR-238 (blue), whereas control samples (no EVs) demonstrated negligible fluorescent signal. **(B)** The RFIs of EVs with miR-39, miR-54, and miR-238 with their corresponding MBs are higher than the different control conditions tested, including EVs with unmatched MBs (a1, a2, a3), EVs from human serum (a4), and no EVs (a5) (n = 3, error bars indicate the standard deviation). **(C)** The RFI for the detection of miR-39 in the engineered EVs increased with increasing concentrations of the cel-miR-39 plasmid transfected into the cells. EV concentrations were held constant at 1.0E^9^ particles/mL for all conditions (n = 3, error bars indicate the standard deviation). **(D)** The ^siEV^PRA was compared against a standard PCR test for detecting miR-39 from engineered EVs generated from transfected Gli36 cells with a plasmid concentration of 400 μg/uL (n = 3, error bars indicate the standard deviation). Representative images and statistical distributions are provided in **Supplementary Fig. 5**. **(E)** The ^siEV^PRA is compared against a standard ELISA for detecting EGFR from EVs isolated from Gli36 cells (n = 3, error bars indicate the standard deviation). Representative images and statistical distributions are provided in **Supplementary Fig. 6**.

The sensitivity of the ^siEV^PRA for RNA detection in siEVs was compared to conventional bulk PCR. EVs harvested from Gli36 cells loaded with 400 ng/mL of the cel-miR-39 plasmid were diluted serially and detected with the ^siEV^PRA and PCR for miR-39.

Signal from the sample was detected at a concentration of 1.0E^6^ vesicles/mL with the ^siEV^PRA (ANOVA, *p* = 0.01), outperforming the detection of miR-39 with PCR, which was undetectable below a concentration of 1.0E^9^ vesicles/mL (**Fig. 2D**, **Supplementary Fig. 5**). Moreover, the sensitivity of the ^siEV^PRA for protein detection in siEVs was compared to standard bulk ELISA. Gli36-derived EVs were diluted serially and detected with both methods for EGFR, a protein upregulated in GBM^6^. Signal from the sample was detected at a concentration of 1.0E^6^ vesicles/mL with the ^siEV^PRA (ANOVA, *p* = 0.01), whereas ELISA could not detect EGFR below a concentration of 1.0E^9^ vesicles/mL (**Fig. 2E**, **Supplementary Fig. 6**). Thus, the ability of the ^siEV^PRA to outperform conventional bulk-analysis methods to detect vesicular RNA and protein, while preserving siEV integrity indicate its potential for the molecular characterization of EV heterogeneity at minimal concentrations.

### 2.3. Simultaneous detection of various biomolecule species

To first determine the ability of the ^siEV^PRA to multiplex various probes at the single-vesicle level, different regions of an mRNA were detected simultaneously within siEVs. Given the length of mRNA strands, three MBs targeting three different regions of the AXL receptor tyrosine kinase (AXL) mRNA, an abundant mRNA found in GBM^27^, were designed such that each MB emitted a different fluorescent signal when hybridized. All three regions of the AXL mRNA were detected in siEVs as single fluorescent spots. Furthermore, magenta, cyan, and yellow spots illustrated the colocalization of two detection probes, whereas white spots demonstrated the colocalization of all detection probes (**Supplementary Fig. 7A**). The probability distributions of the single AXL regions detected by the MBs were similar (**Supplementary Fig. 7B**). Moreover, the fluorescent intensities of the three different regions on the AXL mRNA showed negligibly different fluorescent intensities (ANOVA, *p* = 0.95), demonstrating a uniform and noncompetitive affinity of the MBs to the different regions of the mRNA strand (**Supplementary Fig. 7C**). The colocalization efficiencies for AXL-1 and AXL-2, AXL-2 and AXL-3, AXL-1 and AXL-3, and all three regions were 26.15 ± 2.09 %, 28.31 ± 1.59 %, 22.84 ± 2.52 %, and 3.12 ± 0.58 %, respectively (**Supplementary Fig. 7D**). Given the homogeneity of the MB hybridization to the different regions of the AXL mRNA, the statistically equal colocalization efficiencies for the simultaneous hybridization of two regions (ANOVA, *p* = 0.69) implies a similar probability for two probes to co-detect RNA within siEVs. Furthermore, the fluorophores were only excited when matched by their corresponding emission (**Supplementary Fig. 8**), ensuring the validity of the colocalization as originating from the EV-detecting probes.

To further test the ability of the ^siEV^PRA for multiplexed biomarker detection in siEVs, several combinations of proteins and RNAs were screened. CD63, CD81, and CD9 were detected on Gli36-derived EVs (**Fig. 3A)**. The colocalization efficiencies for CD63 and CD9, CD81 and CD9, CD63 and CD81, and all three proteins were 20.08 ± 2.09 %, 19.31 ± 1.59 %, 20.84 ± 2.52 %, and 2.16 ± 0.58 %, respectively (**Fig. 3B**). EVs harvested from Gli36 cells transfected with cel-miR-39, cel-miR-54, and cel-miR-238 plasmids were detected by their respective MBs (**Supplementary Fig. 9A)**. The colocalization efficiencies for miR-39 and miR-54, miR-54 and miR-238, miR-238 and miR-39, and all miRNA were 32.94 ± 1.47 %, 31.10 ± 1.03 %, 31.26 ± 2.90 %, and 5.51 ± 0.51 %, respectively (**Supplementary Fig. 9B)**. Moreover, proteins and RNAs across various species were detected with the ^siEV^PRA. First, miR-21, miR-9-5p, and AXL were detected in single Gli36-derived EVs to detect different RNA species, including miRNA and mRNA (**Fig. 3A**). The colocalization efficiencies for AXL and miR-9-5p, miR-21 and miR-9-5p, AXL and miR-21, and all three RNA biomarkers were 21.15 ± 2.29 %, 22.62 ± 1.08 %, 20.67 ± 2.58 %, and 2.95 ± 0.18 %, respectively (**Fig. 3B**). Second, CD63, miR-21, and miR-9-5p were detected in single Gli36-derived EVs to multiplex proteins and RNA simultaneously (**Fig. 3A**). The colocalization efficiencies for CD63 and miR-9-5p, miR-21 and miR-9-5p, miR-21 and CD63, and all three biomarkers were 19.30 ± 1.05 %, 22.52 ± 1.90 %, 20.71 ± 2.23 %, and 2.12 ± 0.48 %, respectively (**Fig. 3B**).

**Figure 3:**
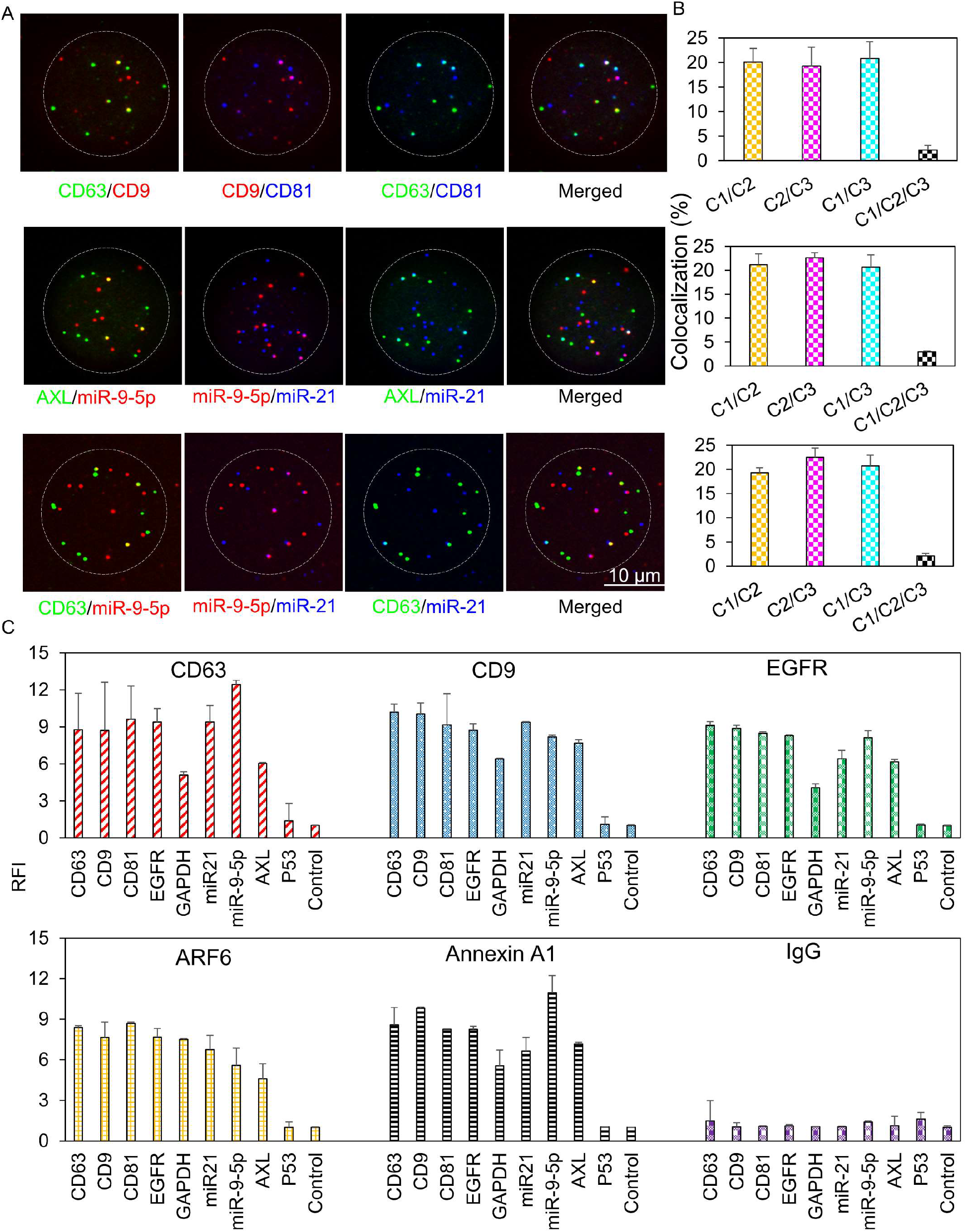
Simultaneous detection across various biomolecules. **(A)** EVs isolated from Gli36 cells are tested with the ^siEV^PRA for three different probes, including multiprotein detection (CD63, CD9, and CD81), mRNA-miRNA detection (AXL, miR-9-5p, and miR-21), and protein-miRNA detection (CD63, miR-9-5p, and miR-21). **(B)** Quantification of colocalization efficiencies for the different biomarkers in Gli36-derived siEVs. C1, C2, and C3 represent the detected biomarkers from left to right (n = 3, error bars indicate the standard deviation). (**C**) The RFIs of CD63, CD9, CD81, EGFR, GAPDH, miR-21, miR-9-5p, AXL, P53, and their corresponding controls, captured by CD63, CD9, EGFR, ARF6, Annexin A1, demonstrates a higher signal for the samples. EVs captured by IgG demonstrate similar signals among the biomarkers as the control (n = 3, error bars indicate the standard deviation).

The ^siEV^PRA platform also enabled the sorting and characterization of siEV subpopulations based on different surface proteins by using different antibodies to capture siEVs and subsequently measure their protein and RNA content. Tetraspanins, ARF6, and Annexin A1 are well-known surface proteins expressed in EVs^28–30^. Nine different biomarkers, including four proteins and five different RNAs, were quantified and revealed different expression levels in siEVs by TIRFM. Fluorescent signals from siEVs showed that expression levels for CD81, CD63, CD9, and EGFR were higher than the other biomarkers independent of the EV subpopulation analyzed. More variability was observed for the different RNAs tested for which there was no clear trend in the level of expression based on the subpopulation analyzed. Thus, these results confirm the heterogeneity in the different EV subpopulations captured on the device (**Fig. 3C**).

### 2.4. Single-EV analysis of RNA biomarkers in EV subpopulations and clinical samples

To validate the use of the ^siEV^PRA for characterizing EV heterogeneity, we performed transcriptomic analysis of six different GBM cell lines and their corresponding EVs, including SF268, SF295, SF539, SNB19, SNB75, and U251 using microarray and small RNA sequencing. We found several RNAs that exhibited high concentrations in cells and EVs (**Fig. 4A**). Among them, four transcripts, two mRNAs (NSF and NCAN) and two miRNAs (miR-9-5p and miR-1246-5p) were selected for further analysis, since they have also been reported to be associated with GBM^31–34^. The concentrations of the four selected transcripts measured in the different GBM cell lines showed less variability than their corresponding EVs (**Fig. 4B**). The heterogeneity of these transcripts in EVs was further explored with the ^siEV^PRA. NSF, NCAN, miR-9-5p, and miR-1246-5p were measured in siEVs (**Fig. 4C, Supplementary Fig. 10-13**). siEVs from the six cell lines exhibited higher RFIs across the four biomarkers than the control samples (ANOVA, *p* < 0.0001). Although the four RNAs were detectable in siEVs, variations in fluorescent signal across EVs from the different cell lines demonstrated vesicular heterogeneity. The statistical distributions for siEV intensity illustrate a more homogeneous expression for the mRNAs compared to the miRNAs. For NSF and NCAN, the distribution maxima were relatively consistent across the EVs from the six cell lines at 356.09 ± 64.20 and 300.14 ± 78.02, respectively (**Supplementary Fig. 10 & 11**). On the other hand, miR-9-5p from SF268-, SF295-, SF539-, and SNB75-derived siEVs had more heterogeneous profiles with distribution maxima shifted to the right; similarly, miR-1246-5p from SF268-, SNB75-, and SNB19-derived siEVs also demonstrated a heterogeneous expression with distribution maxima shifted to the right (**Supplementary Fig. 12**). In general, distribution maxima for the miRNA had more variability. Specifically, miR-1246-5p showed more significant discrepancies than other RNAs (**Supplementary Fig. 13**). Distribution maxima among all controls showed less variability with values at 149.15 ± 28.92.

**Figure 4:**
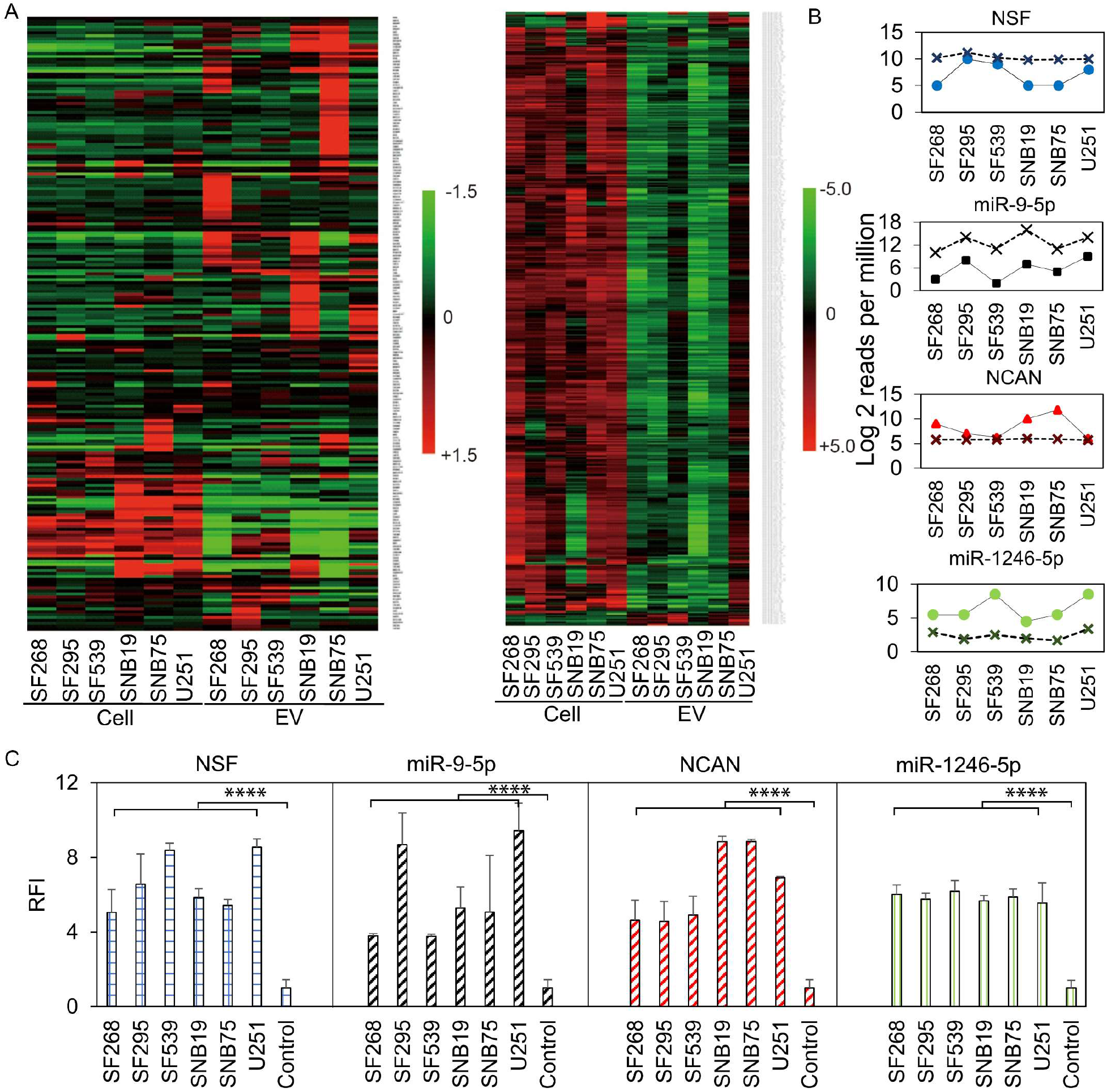
RNA sequencing of GBM-specific biomarkers and validation at a singlevesicle resolution. **(A)** Cellular and vesicular mRNA (left) and miRNA (right) are sequenced across six GBM cell lines, including SF268, SF295, SF539, SNB19, SNB75, and U251, revealing the upregulation of NSF, miR-9-5p, NCAN, miR-1246-5p in cells and EVs. **(B)** NSF, miR-9-5p, NCAN, and miR-1246-5p are profiled in EVs (solid line) and cells (dash line) from the six different GBM cell lines by NGS bulk characterization (n = 3). **(C)** NSF, miR-9-5p, NCAN, and miR-1246-5p are profiled in siEVs from the six different GBM cell lines with the ^siEV^PRA, showing upregulation of the four RNA biomarkers in comparison to the control samples (n = 3, error bars indicate the standard deviation).

Finally, the ^siEV^PRA was used to characterize EV subpopulations from GBM patient serum. An average of 20 μL of purified serum was processed from GBM patients at different stages of treatment (*n* = 10). Serum from healthy individuals was also processed as healthy controls (*n* = 10). For the GBM patient samples, we measured fluorescent signals for NSF, NCAN, miR-9-5p, and miR-1246-5p RNAs in siEVs, while fluorescent signals in EVs from healthy donor serum were significantly lower (*p* < 0.0001, **Fig. 5**). Comparisons of the statistical distributions of siEV intensity for the different RNA species revealed that NSF and NCAN mRNAs presented more homogeneous fluorescent signals with distribution maxima at 355.80 ± 2.76 and 383.47 ± 28.92,. On the other hand, the statistical distributions for miR-9-5p and miR-1246-5p miRNAs showed a broad distribution of fluorescent signals with distribution maxima at 1482.67 ± 32.16 and 1136.06 ± 27.43, respectively. However, the statistical distributions of the different RNAs measured from healthy donor serum exhibited less variability, with distribution maxima at 210.16 ± 32.76. Similar trends were observed with the control samples (**Supplementary Fig. 14**). These findings confirm the ability of the ^siEV^PRA to measure RNA heterogeneity in siEVs from complex biofluids. The success of this work opens the possibility for its application in liquid biopsies for cancer diagnoses and prognoses.

**Figure 5.**
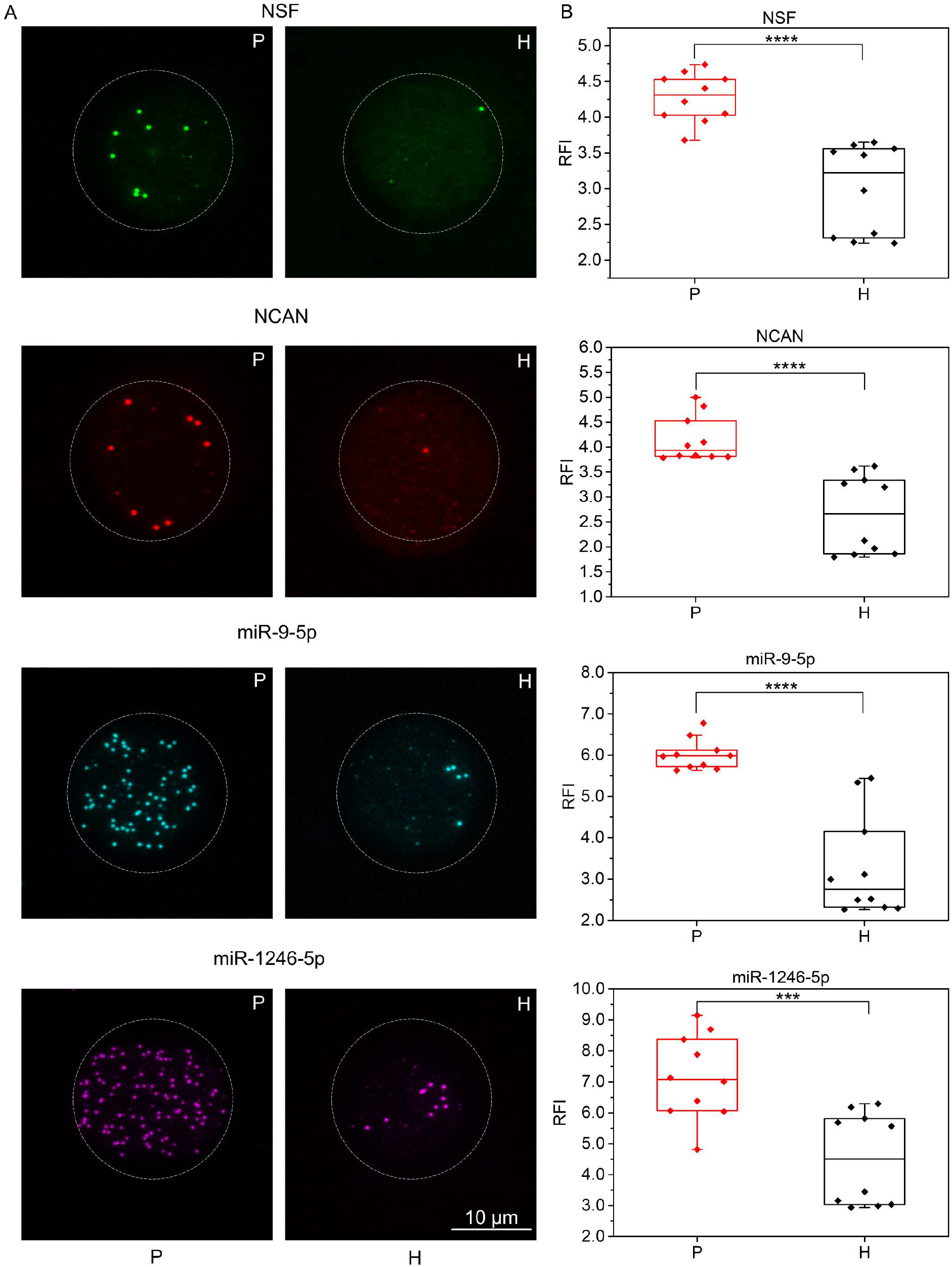
Measurements of GBM-specific biomarkers at a single-vesicle resolution from GBM patient serum. **(A)** Representative TIRFM images of siEV NSF, NCAN, miR-9-5p, and miR-1246-5p biomarkers from GBM patients (P) (n = 10) and healthy donor (H) control serum (n = 10) characterized with the ^siEV^PRA. **(B)** The RFI signals of NSF, miR-9-5p, NCAN, and miR-1246-5p in the GBM sample are higher than the RFI signals obtained for the healthy donor and control samples (n = 10).

## 3. Conclusion

The physical and biological heterogeneity that EVs exhibit has made their accurate molecular quantification a difficult task^4,6^. Current methods for the molecular analysis of EVs, including WB, ELISA, PCR, NGS, and MS, require a high concentration of EVs and a breakdown of their vesicular structure to access their cargo. As a result, crucial molecular information of tissue-specific or disease-specific EVs is lost^35^. To overcome these limitations, several new technologies have been developed to isolate and characterize EVs *in situ*^36–40^. However, these platforms still require many EVs per measurement and are limited in characterizing different types of molecular cargo. We developed the ^siEV^PRA as a promising technology that enables the robust investigation of heterogeneity in siEV cargo. The ^siEV^PRA was built as an array of microdomains on a polymer-coated glass surface fabricated by maskless UV photopatterning. Antibodies immobilized within the arrayed surface targeted siEV subpopulations that were detected *in situ* with fluorescently labeled antibodies and RNA-targeting MBs. 20 μL of complex biofluids (e.g., cell culture media (CCM), serum) is enough to perform a multiparametric characterization of various proteins and RNAs in siEVs to investigate vesicular heterogeneity.

The higher sensitivity of the ^siEV^PRA versus traditional bulk-analysis methods offers an alternative assay for analyzing biomarker heterogeneity in EVs. An advanced engineered EV model system was used^41^ to test the ability of the ^siEV^PRA to measure differences in vesicular RNA. Three non-human miRNAs, including miR-39, miR-54, and miR-238 were engineered in EVs whereby their heterogeneity was detected. Similarly, the analysis of tetraspanin co-expression, which are abundant protein biomarkers in EVs^42^, was analyzed on siEVs demonstrating low colocalization efficiencies, which agrees with reported siEV tetraspanin analyses^43^. Although tetraspanins are highly expressed on EVs implying high co-expression, our siEV analysis demonstrated a heterogeneous expression of different tetraspanins not detectable with bulk characterization methods. Interestingly, when multiplexing across various biomolecule species, the colocalization efficiency was lower for protein-RNA detection compared to RNA-RNA and protein-protein detection. The difference may be attributed to the location of the biomolecules, since RNAs exist within the aqueous core of the EV, whereas tetraspanins are typically localized on the EV membrane surface^44^. Regardless, the successful multiplexing of protein and RNA by the ^siEV^PRA can expand the field of EV heterogeneity analyses.

With the ^siEV^PRA, subpopulations of EVs from GBM cell lines demonstrated vesicular heterogeneity in protein, mRNA, and miRNA expression through colocalization analyses and were validated by bulk RNA sequencing. A comparative molecular analysis between RNA sequencing and the ^siEV^PRA showed the possibility of integrating workflows for the discovery and validation of disease-specific RNA biomarkers, especially for RNA species enriched in EVs. Bulk RNA sequencing of GBM cell lines and their corresponding EVs revealed a subset of RNAs present at different concentrations among EVs and their parental cells. Some of the RNAs analyzed exhibited higher concentrations within EVs, which may be a result of selective packing or novel EV subtypes. This further emphasizes the importance of understanding the complex and diverse biogenesis of EVs and the mechanisms of molecular packing. Interestingly, for the miRNAs measured, a similar trend in the concentrations was observed between bulk sequencing and the ^siEV^PRA; however, some discrepancies were observed between microarrays and the ^siEV^PRA for mRNA measurement. Given that mRNAs are long, single-stranded RNA molecules, whereas miRNAs are small single-stranded non-coding molecules^45,46^, miRNA may hybridize more efficiently with the MBs as they have similar base-pair lengths. Perhaps generating MBs targeting different regions of the mRNA as was done for AXL would improve the consistency between bulk and siEV mRNA analyses.

The feasibility of analyzing RNAs in siEVs from GBM patient serum samples unveils the potential of the ^siEV^PRA technology for various liquid biopsy applications with unmatched levels of sensitivity. Applying GBM patient serum to the ^siEV^PRA, different mRNAs and miRNAs associated with GBM were validated and demonstrated the assay’s clinical potential. Although the current study focused on cancer biomarker analysis, the ^siEV^PRA can be easily adapted to other diseases. Furthermore, changing the capture antibodies within the microdomains can be used to sort EVs based on subpopulations, which may uncover differences in subpopulation-dependent packing of biomolecules and illuminate biogenesis pathways that conventional bulk-analysis methods may muddle. Lastly, the ability for the ^siEV^PRA to multiplex across various biomolecule species offers a unique opportunity to study EV heterogeneity more comprehensively than has been previously accomplished.

## 4. Experimental Section

### Materials

0.01 % (w/v) poly-L-lysine (PLL, MilliporeSigma, Burlington, MA), 5 kDa methoxy-poly(ethylene glycol)-succinimidyl valerate (mPEG-SVA) (ThermoFisher Scientific, Waltham, MA), 0.1 M 4-(2-hydroxyethyl)-1-piperazineethanesulfonic acid (HEPES) buffer (pH = 8.5) (ThermoFisher Scientific, Waltham, MA), 4-benzoylbenzyl-trimethylammonium chloride (PLPP) (Alvéole, France), NeutrAvidin (NA) (ThermoFisher Scientific, Waltham, MA), bovine serum albumin (BSA) (Millipore Sigma, Burlington, MA),

Tris-Ethylenediaminetetraacetic acid (EDTA) buffer (ThermoFisher Scientific, Waltham, MA), *E. coli* (VB200815-1011zys), *E. coli* (VB200815-1012qpx), *E. coli* (VB200815-1013ugb) (VectorBuilder Inc., Chicago, IL). All capture and detection antibodies used in the study are provided in **Supplementary Table 2**. Capture antibodies were biotinylated using an EZ-Link^™^ micro Sulfo-NHS-biotinylation kit (ThermoFisher Scientific, Waltham, MA). All RNA molecular beacons (MBs) used in the study are provided in **Supplementary Table 3**.

### Substrate fabrication

Coverslips were cleaned with ethanol and then deionized (DI) water via sonication for 3 min. The surface of the coverslip was treated with oxygen plasma for 1 min to activate the surface. A small drop of 0.01 % (w/v) PLL was placed onto parafilm, where the treated coverslip was then placed for an even distribution of the PLL. After incubating the coverslip for 30 min at room temperature (RT), the PLL-coated coverslip was rinsed with DI water and dried with nitrogen flow. Following the same method, 100 mg/mL of mPEG-SVA diluted in 0.1 M HEPES was evenly distributed on the PLL-coated coverslip. The coverslip was incubated at RT for 1 h before rinsing with DI water and drying with a nitrogen airflow. The treated coverslip could be stored for three weeks at 4 °C before use^47^.

### Device fabrication and surface modification

The passivated coverslip was photopatterned using the PRIMO optical module (Alvéole, France) mounted on an automated inverted microscope (Nikon Eclipse Ti Inverted Microscope System, Melville, NY). Briefly, grayscale images were translated into UV light via a DMD that allows for a maskless illumination of different UV intensities correlating to the corresponding grayscale values^21^. Following the passivation of the coverslip, PLPP gel was diluted in 96 % ethanol to distribute the gel throughout the surface of the coverslip evenly. After the ethanol evaporated, a silicone spacer (W x L 3.5 mm x 3.5 mm, 64 wells, Grace Bio Labs, Bend, OR) was placed on top of the PEG-coated coverslip. A five-by-five array of 20-μm diameter circles spaced 80 μm center-to-center was exposed onto the coverslip with the PRIMO optical module. To optimize the relative fluorescence intensity (RFI) between samples and their controls of the detection probes, microdomains at different grayscale values, including 0, 25, 50, 75, and 95 % with UV doses, including 10, 20, and 30 mJ/mm^2^ were examined (**Supplementary Table 1**). After the UV illumination was completed, the photoetched coverslip was washed under a stream of DI water and dried by nitrogen flow. A microscopy slide (Fisher Scientific) was placed under the coverslip, and the 64-well ProPlate microarray system (Grace Bio Labs) was placed gently on the faced-up photoetched coverslip. The assembled array was secured by self-cut Delrin snap clips (Grace Bio Labs) to avoid leakage or potential contamination. The photoetched coverslip was rehydrated with phosphate-buffered saline (PBS) for 15 min before antibody functionalizing of the microdomains.

### Cell culture

U251 and Gli36 GBM cell lines were cultured in Dulbecco’s Modified Eagle Medium (DMEM). SF268, SF295, SF539, SNB19, and SNB-75 GBM cell lines were cultured in Roswell Park Memorial Institute (RPMI) 1640 medium. All cell culture media was prepared with 10 % (v/v) fetal bovine serum (FBS) and 1 % (v/v) penicillinstreptomycin. Cell lines were first cultured to 90 % confluence at 37 °C in a 5 % CO_2_ incubator. Before EV collection, cells were washed with PBS twice, after which the cells were incubated in media supplemented with 10 % (v/v) EV-depleted FBS. The FBS was filtered by tangential flow filtration (TFF) (MWCO: 300 kDa) from which the permeate containing EV-depleted FBS was used^48^. After two days of cell culture, the EV-enriched cell culture medium (CCM) was collected and centrifuged at 2,000 x *g* for 7 min at RT to separate cell debris before further analysis.

### Human tumor specimen collection

Human tumor tissue was obtained under Institutional Review Board (IRB)-approved protocols at MD Anderson Cancer Center (PA 19-0661) in accordance with national guidelines. All patients signed informed consent forms during clinical visits before surgery and sample collection. Patients did not receive compensation in return for their participation in this study.

### Engineered-EV RNA model system

Cell transfection was conducted via a cellular nanoporation (CNP) biochip^41^. Briefly, a single layer of Gli36 cells (~ eight million) was spread overnight on a 1 cm × 1 cm 3D CNP silicon chip surface. Cel-miR-39, cel-miR-54, and cel-miR-238 plasmids at a weight ratio of 1:1:1 were pre-mixed at a concentration of 100 ng/mL each in PBS for transfection. The plasmid solution was injected into individual cells via nanochannels using a 200 V electric field for a total of 5 pulses, at 10 ms durations and 0.1 s intervals. EVs were collected from the cell supernatant after 24 h of the cell transfection.

### Healthy donor serum collection

10 mL of whole blood from healthy donors was collected into BD Serum Separation Tubes (SST) (Thermo Fisher Scientific, Waltham, MA). SSTs were gently placed upright to coagulate for 60 min after being rocked 10 times. The SSTs were centrifuged at RT at 1,100 × *g* for 10 min. The serum was stored in 1 mL aliquots at −80 °C. All blood samples were collected under an approved Institutional Review Board (IRB) at The Ohio State University (IRB #2018H0268).

### EV purification

The EV-enriched CCM and healthy donor serum were introduced into a TFF system as described by our previous technique to purify EVs^48^. In brief, CCM or serum was circulated through a 500 kDa TFF hollow fiber filter cartridge, where EVs were retained and enriched in the system (~ 2 mL), while free proteins and nucleic acids permeated through the filter. Further diafiltration cycles with PBS were performed until pure EVs were obtained (150 mL of PBS in ~ 80 min). The EVs were further enriched by spinning down the sample within a 10 kDa ultracentrifugal unit at 3000 × *g* at 4 °C until a final volume of 100 μL was achieved.

### EV size and concentration quantification

A tunable resistive pulse sensing (TRPS) method, qNano Gold (Izon Sciences, Boston, MA), was employed to quantify the size and concentration of EVs^49^. 35 μL of the sample was pipetted into NP100 (50 – 330 nm) and NP600 (275 – 1570 nm) nanopore membranes. A pressure of 10 mbar and a voltage of 0.48 and 0.26 V was applied for the NP100 and the NP600, respectively. Polystyrene nanoparticles (CPC100 and CPC400) were used to calibrate the samples.

### MB design and quantification

MBs (listed 5–3’) targeting RNAs detected in this study are provided in **Supplementary Table 2**. Locked nucleic acid (LNA) nucleotides (depicted as +) were incorporated into oligonucleotide strands to improve the thermal stability and nuclease resistance of the MBs for incubation at 37 °C. The designed MBs were custom synthesized and purified using high-performance liquid chromatography (HPLC) (Integrated DNA Technologies, Coralville, IA).

### EV protein and RNA staining

10 μg/μL of MBs diluted in 1X Tris-EDTA (TE) buffer were mixed with the EV sample for 1 h at 37 °C. As for protein detection, 0.4 μg/mL of fluorescently labeled monoclonal antibodies were diluted into a solution of 3 % BSA and 0.05 % (v/v) Tween^®^ 20 in PBS and were added to the EV sample for 1 h at RT. For single biomarker analysis, sole detection probes were added. To analyze multiple proteins or RNAs, the probes were added sequentially, monoclonal antibodies were added first, followed by MBs.

### Single EV capture using the ^siEV^PRA

0.1 mg/mL of NA was added to the chip and allowed to physisorb onto the photocleaved microdomains for 30 min. The chip was washed with PBS thoroughly to remove excess NA. A blocking solution of 3 % BSA and 100 mg/mL of mPEG-SVA was added to avoid unwanted non-specific binding. Subsequently, biotinylated anti-CD63 and anti-CD9, anti-EGFR, anti-ARF6, anti-Annexin A1, and anti-IgG were added at 20 μg/mL each and allowed to sit overnight at 4 °C. 3 % BSA was added for 1 h to further block after the capture antibodies were washed away. EVs were then added and allowed to tether to the antibodies for 2 h at RT. Unbounded EVs were later washed away with PBS.

### Image analysis

The images of fluorescently labeled siEVs were obtained by TIRFM (Nikon Eclipse Ti Inverted Microscope System, Melville, NY) with a 100× oil immersion lens. An automatic algorithm quantified the TIRFM images by detecting all bright spots by determining the outline of each bright spot as defined by varying fluorescent intensities throughout the image. The background noise was removed using a Wavelet de-noising method, and each bright spot’s net signal was obtained. The sum of all the bright spots within each microdomain was employed to calculate the total fluorescence intensity of the sample alongside a statistical distribution of the mean fluorescent intensity. The total fluorescence intensity of samples was normalized to the total fluorescence intensity of negative controls as relative fluorescence intensities (RFI)^53^.

### Enzyme-linked immunosorbent assay (ELISA)

Epidermal growth factor receptor (EGFR) protein expression levels on the surface of Gli36-derived EVs were quantified using an EGFR Human ELISA kit (ThermoFisher Scientific, Waltham, MA). EVs were spiked in healthy donor serum at different concentrations ranging from 0 to 1.0E^11^ particles/mL while maintaining the serum-derived EV concentration at 1.0E^9^ particles/mL. EGFR expression was quantified according to the manufacturer’s instructions.

### qRT-PCR

Cel-miR-39-3p levels within the engineered EVs were quantified using qRT-PCR. Total RNA from the cells and EVs was isolated and purified using an RNeasy Mini Kit and a miRNeasy Serum/Plasma kit (Qiagen, Hilden, Germany), respectively, according to the manufacturer’s instructions. cDNA was synthesized from the total RNA using a High-Capacity cDNA Reverse Transcription kit (Applied Biosystems, Foster City, CA) on a thermal cycler (Veriti 96-Well Thermal Cycler, Applied Biosystems). Cel-miR-39-3p expression was quantified using a TaqMan Gene Expression assay (ThermoFisher Scientific, Assay Id: Hs01125301_m1) on a Real-Time PCR System (Applied Biosystems, ThermoFisher Scientific).

### Scanning electron microscopy (SEM)

TFF-purified EVs were tethered to the micropatterned coverslip overnight at 4°C. The tethered EVs were fixed in a 2 % glutaraldehyde (MilliporeSigma, Burlington, MA) and 0.1 M sodium cacodylate solution (Electron Microscopy Sciences, Hatfield, PA) for 3 hr. EVs were incubated in 1 % osmium tetraoxide (Electron Microscopy Sciences) and 0.1 M sodium cacodylate for 2 h after washing with a 0.1 M sodium cacodylate solution. Subsequently, the sample was dehydrated with increasing ethanol concentrations (50, 70, 85, 95, and 100 %) for 30 min each. Later, the CO_2_ critical point dryer (Tousimis, Rockville, MD) was applied to dry the sample. Last, a ~ 2 nm layer of gold coating was completed using a sputtering machine (Leica EM ACE 600, Buffalo Grove, IL) and was imaged using an SEM (Apreo 2, FEI, ThermoFisher Scientific).

### RNA sequencing

RNA, including miRNA, was isolated from cells and cell-derived EVs using the miRNeasy kit (QIAGEN, Germantown MD). The RNA was eluted with 50 μl of nuclease-free H20, and the quality was assessed using an RNA (Pico) chip on an Agilent 2100 Bioanalyzer (Agilent Technologies, Santa Clara, CA). A small RNA sequencing library construction method that utilizes adapters with four degenerated bases to reduce adapter-RNA ligation bias was used to characterize the miRNA (PMID: 29388143). Size selection was performed using a Pippin HT automated size-selection instrument (Sage Science, Beverly, MA), and library concentrations were measured with the NEBNext Library Quant Kit (New England Biolabs, Ipswich, MA). The libraries were pooled to a final concentration of 2 nM and run on a NextSeq sequencer (Illumina, San Diego, CA). The small RNA sequencing (sRNA-Seq) data was analyzed with sRNAnalyzer^50^. The quantity of miRNA was determined based on the number of mapped reads that were normalized with Count Per Mapped Million (CPM). RNA from cells and EVs were analyzed using Agilent Human Whole Genome 8×60 microarrays with fluorescent probes prepared from isolated RNA samples using Agilent QuickAmp Labeling Kit according to the manufacturer’s instructions (Santa Clara, CA). Gene expression information was obtained with Agilent’s Feature Extractor and processed with the in-house SLIM pipeline^51^.

### Colocalization efficiency

An open-source plugin for ImageJ called EzColocalization was employed to visualize and measure the colocalization of EV biomarkers from acquired TIRFM images^52^.

### Statistical analysis

Data are expressed as the mean ± SD. A significant test between different mean values was evaluated using the JMP Pro 14 software (JMP, Cary, NC). Differences between samples were considered statistically significant for p < 0.05.

## Supporting information

Supporting information is available from the journal website or the authors.

## Acknowledgment

We acknowledge all cancer patients and healthy volunteers who participated in this study. This work was supported by the U.S. National Institutes of Health (NIH) grants UG3/UH3TR002884 and U18TR003807 (ER). Additional support for ER was provided by the William G. Lowrie Department of Chemical and Biomolecular Engineering and the Comprehensive Cancer Center at The Ohio State University.

## Contributions

ER conceived the idea. ER, JZ, and LTHN designed the study. JZ, XW, and XYR performed experiments and data analysis. ER, JZ, XW, XYR, LTHN, and NW prepared all tables and figures and drafted the original manuscript with input from all authors. XW developed the quantification codes. LTHN established protocols for protein/RNA analysis. YM performed cell transfection and PCR. NW performed healthy donor serum samples collection. KJK designed molecular beacons for all RNA targets. MY performed EV purification experiments. SF, KS, IL, and KW performed bulk RNA sequencing analysis for cells and EVs. AFP provided supplies and equipment for TFF purification of EVs. KH, DL, YW, JH, WJ, and BYSK coordinated all the efforts with the clinic, obtained required permissions, and selected clinical samples for the study. LJL supervised the design of molecular beacons and edited the manuscript. ER supervised the whole study and edited the manuscript. All authors provided critical feedback and helped to shape the research, analysis, and manuscript.

## Data availability statement

The data supporting the findings of this study are available from the corresponding author upon reasonable request.

