## Supporting Information for "Engineering a Single Extracellular Vesicle Protein and RNA Assay (^siEV^PRA) via In Situ Fluorescence Microscopy in a UV Micropatterned Array"

**Supplementary Table 1: RFI optimization by controlling monolayer degradation.**

| Gray Scale / Dose ($mJ/mm^2$) | 100% | 95% | 75% | 50% | 25% | 0% |
|---|---|---|---|---|---|---|
| 30 | 1 | 1.5 | 4 | 2.5 | 1 | 1 |
| 20 | 1 | 1 | 3.5 | 6 | 1 | 1 |
| 10 | 1 | 1 | 1.5 | 3.5 | 1 | 1 |

**Supplementary Table 2: List of used antibodies.**

| Function | Target | Vendor | Catalog No. |
|---|---|---|---|
| **Capture Antibody** | CD63 | R&D system | MAB5048 |
| | CD9 | R&D system | MAB1880 |
| | EGFR | ImClone LLC | Erbitux-C225 |
| | ARF6 | R&D system | NBP2-41263 |
| | Annexin A1 | R&D system | MAB3770 |
| | IgG | Invitrogen | 31903 |
| **Detection Antibody** | CD63 Alexa Fluor 488 | Santa Cruz Biotechnology | #sc-5275 |
| | CD9 Alexa Fluor 594 | Santa Cruz Biotechnology | #sc-59140 |
| | CD81 Alexa Fluor 640 | Santa Cruz Biotechnology | #sc-166029 |
| | EGFR Alexa Fluor 561 | Cell Signaling | #5616-D38B1 |

**Supplementary Table 3: List of MB designs.**

| RNA | MB |
|---|---|
| miR-21 | T+CA A+CA /iCy3/ +TCA +GT+C T+GA TAA GCT AAC TTA TCA GAC TGA /3BHQ_2/ |
| miR-21 | T+CA A+CA /iCy5/+TC A+GT +CT+G ATA AGC TAA CTT ATC AGA CTG A/3IAbRQSp/ |
| AXL-1 | +CTC CC+C/i6-FAMK/+GG A+TT T+GG CA+C TGC AGT GCCAAA TCC /3BHQ_1/ Product |
| AXL-2 | +GTG +ATT/iCy5/+CT G+AG C+TG G+CT GAC CAA GCCAGC TCA G/3IAbRQSp/ |
| AXL-3 | AXL +TGG +TGT/iCy3/+CT A+GT T+AG T+CA CAA CTG TGT GACTAA CTA G/3BHQ_2/ |
| miR-9-5p | +CAT +ACA /iCy3/+GC T+AG A+TA ACC AAA +GAT TGG TTA TCT AGC /3BHQ_2/ |
| NSF | +GTT +G/iFluorT/C +CCA +CTG +AGA +AGG CCT TCT CAG TGG /3BHQ_1/ |
| NCAN | +GCT +CCA /iCy3/+GG C+AT A+TC C+AC CTC ATG GAT ATG CC/3BHQ_2/ |
| miR-1246-5p | +CCT +GC/iCy5/ +TC+C AA+A AA+T CCA TTC GGA TTT TTG GA/3IAbRQSp/ |
| Cel-miR-39-3p | +CAA +GC/iFluorT/ +GAT +TTA +CAC +CCG GTG AGG GTG TAA ATC /3BHQ_1/ |

| | |
|---|---|
| Cel-miR-54-3p | +GTT +CTC /iCy3/+GT C+GT C+TC A+TA TCC TGG ATA TGA GAC GAC /3BHQ_2/ |
| Cel-miR-238-3p | +CTT T+GA A/iCy5/+C GT+C C+GA +GAA CAT CCA TTC TCG GAC G/3IAbRQSp/ |
| P53 | +CTC CGT /CY5/CAT GTG CTG TGA CTT CAC AGC ACA TG /BHQ3-3/ |

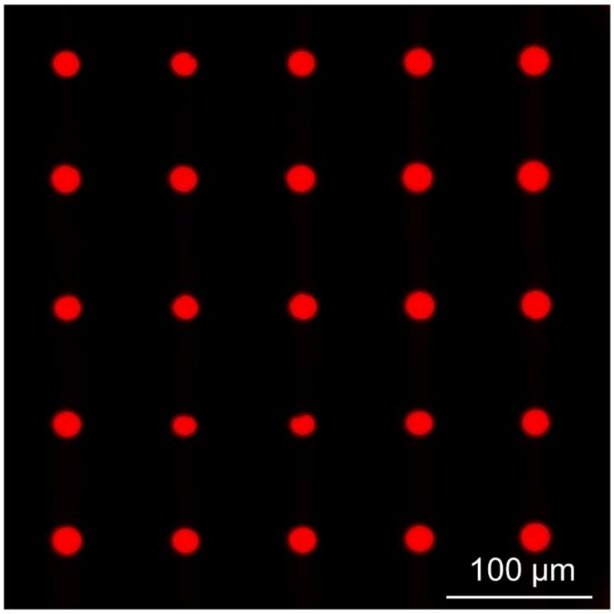


**Supplementary Figure 1: Homogenous NA physisorption.** UV degradation of the PEG monolayer allows the homogenous adsorption of NA in distinct microdomains.

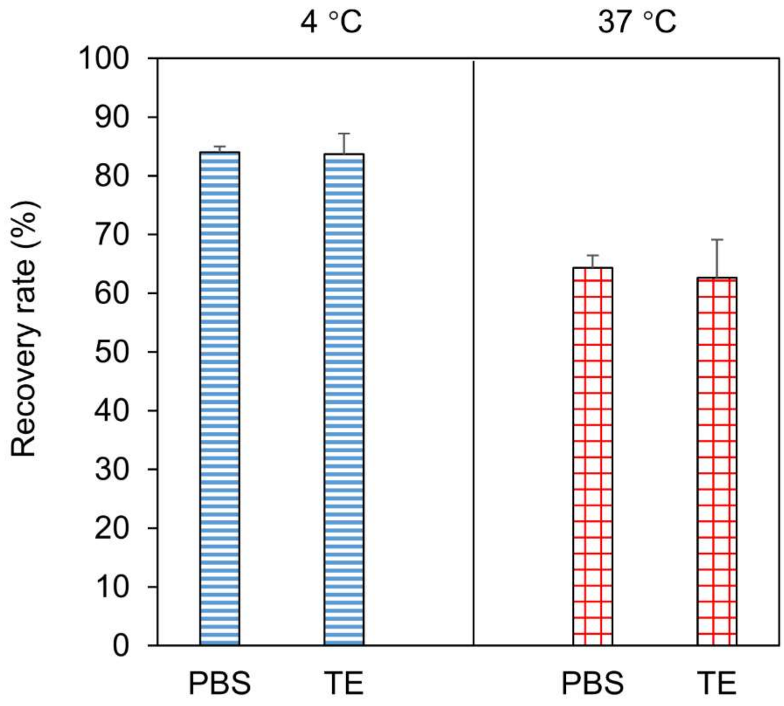


**Supplementary Figure 2: EV recovery rate in different buffers.** The concentration of EVs before and after incubation with a TE buffer and PBS at 4 °C and 37 °C demonstrate equal recovery rates (n = 3, error bars indicate the standard deviation).

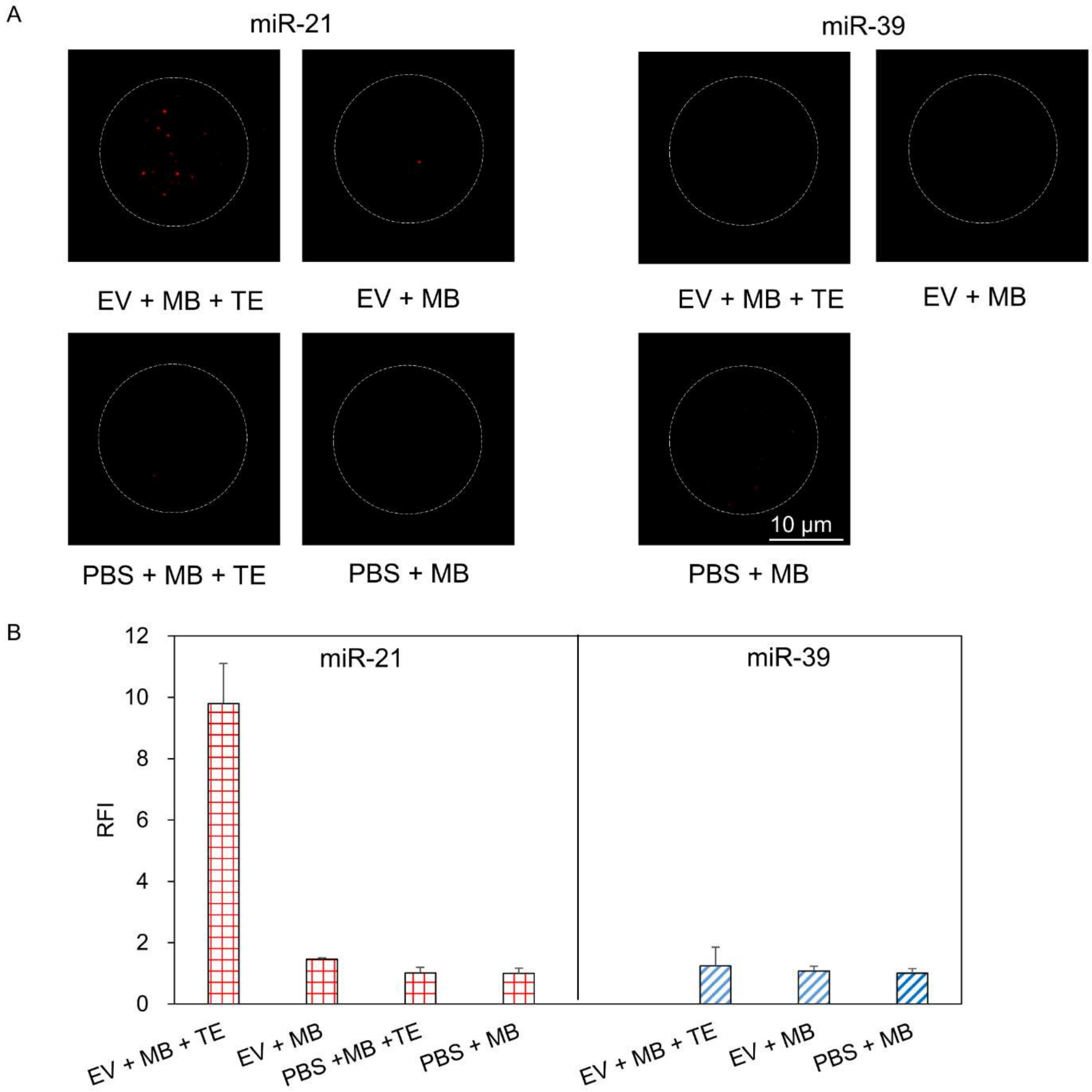


**Supplementary Figure 3: Specificity of RNA detection via MBs**. **(A)** miR-21 and miR-39 are detected with the $^{siEV}$PRA in Gli36-derived siEVs with and without partial permeabilization of the lipid membrane. **(B)** The qualitative images are quantified as RFIs on the Gli36-derived siEVs (n = 3, error bars indicate the standard deviation).

signal.

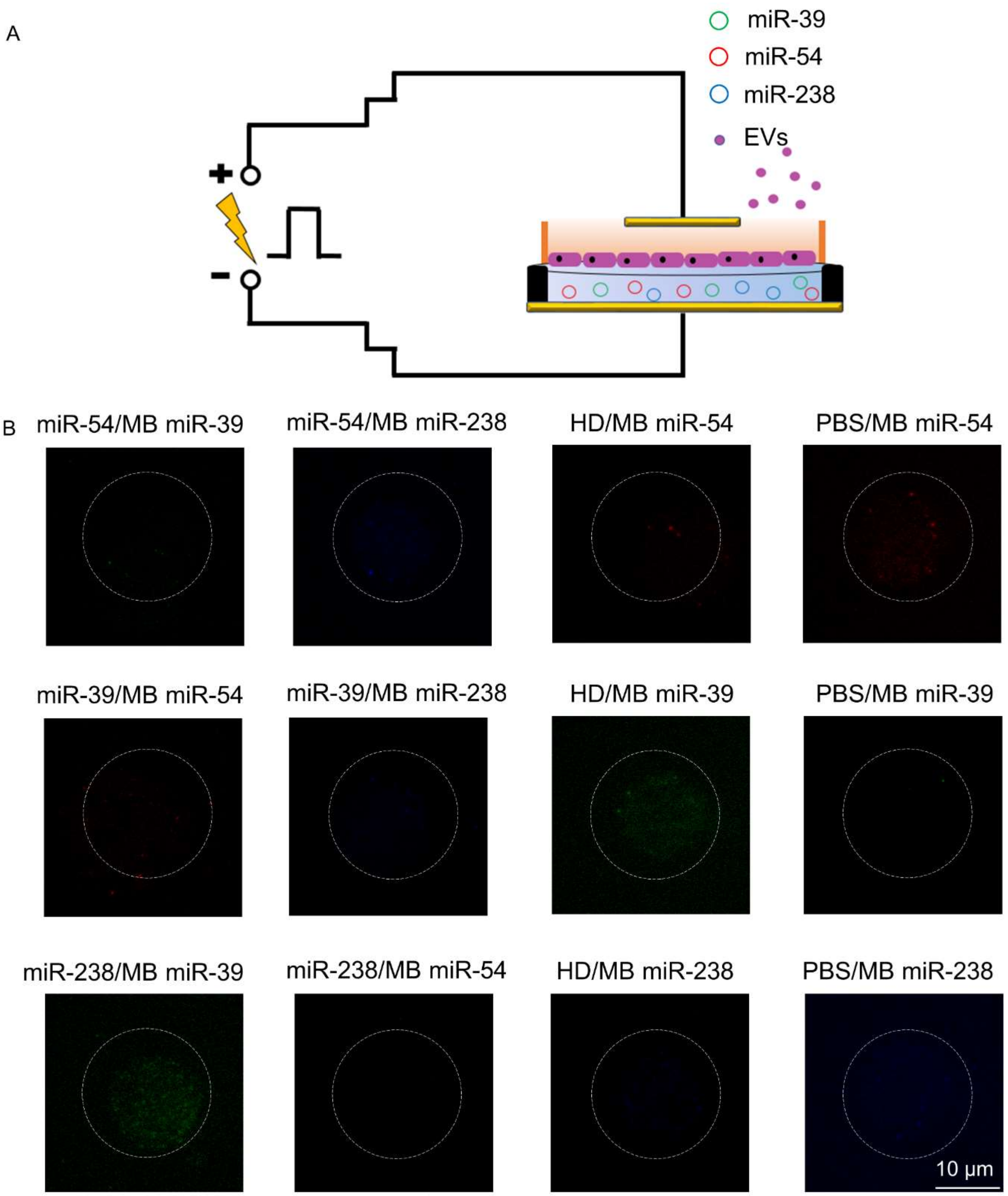


**Supplementary Figure 4: Cross-reactivity of engineered EVs**. **(A)** Gli36-derived EVs were transfected with cel-miR-39, cel-miR-54, and cel-miR-238 with the CNP biochip[41]. **(B)** The unmatched MB controls and the healthy serum control produced a low fluorescent signal.

A

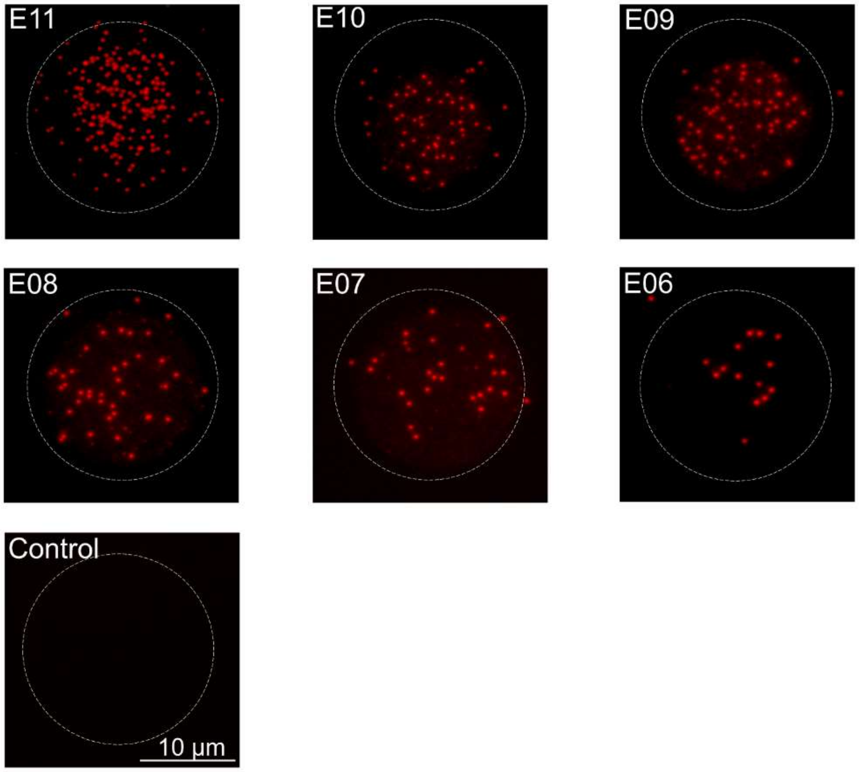
E11
E10
E09
E08
E07
E06
Control
10 μm

B

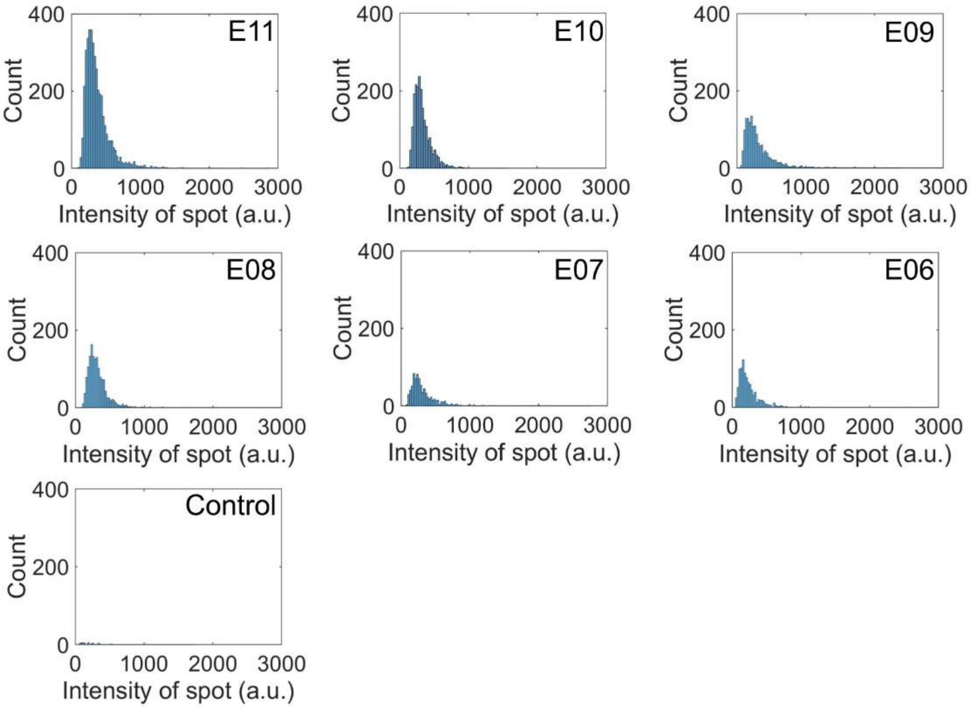
E11
E10
E09
E08
E07
E06
Control
Count
400
200
0
0
1000
2000
3000
Intensity of spot (a.u.)

**Supplementary Figure 5: Sensitivity of the $^{siEV}$PRA for RNA detection. (A)** A serial dilution of engineered EVs enriched with miR-39 was detected with the $^{siEV}$PRA. **(B)** The qualitative images are quantified as statistical distributions.

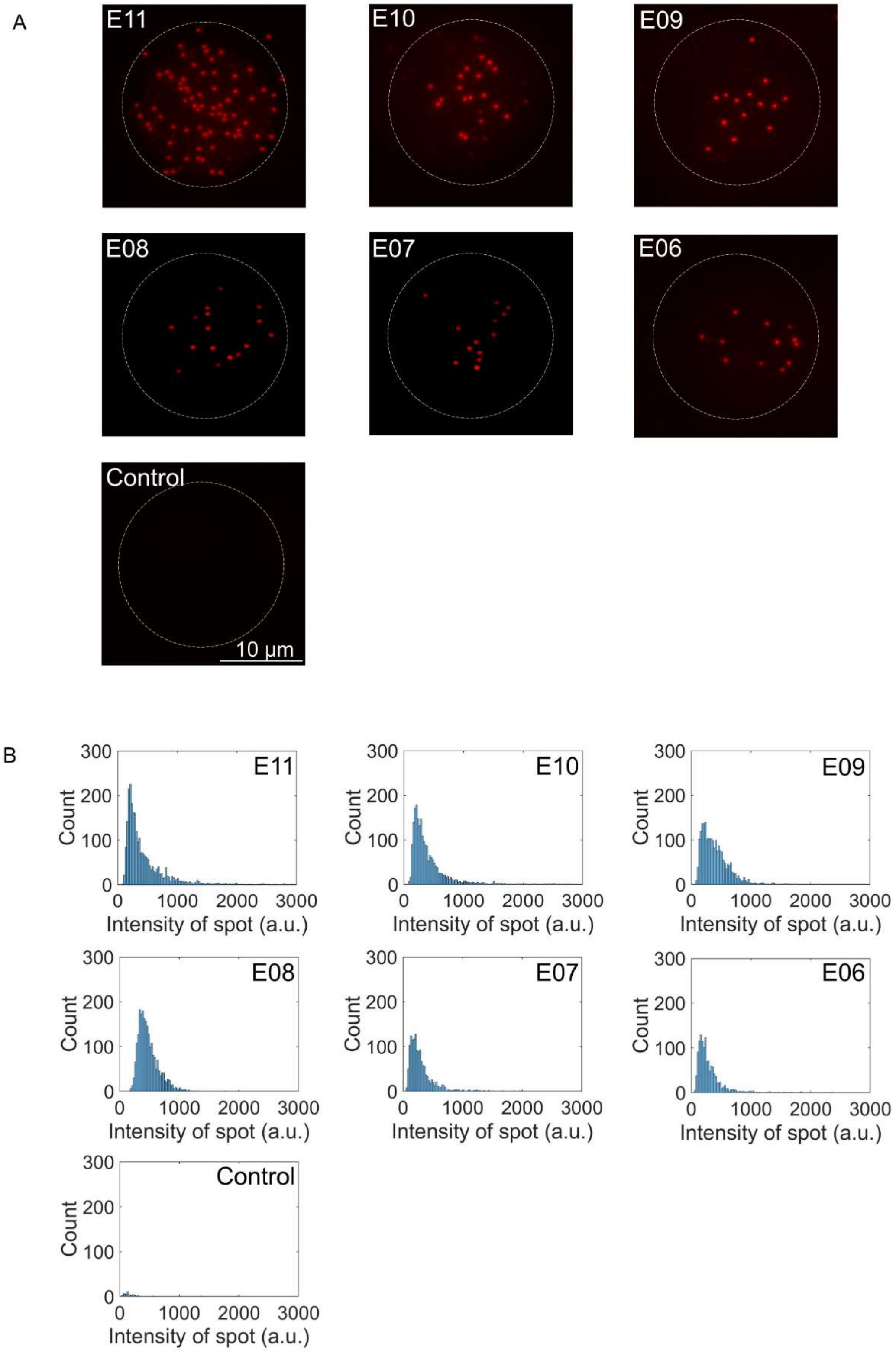
A
E11
E10
E09
E08
E07
E06
Control
10 μm
B
E11
E10
E09
E08
E07
E06
Control
Count
Intensity of spot (a.u.)
0
100
200
300
1000
2000
3000

**Supplementary Figure 6. Sensitivity of the $^{siEV}$PRA for protein detection. (A)** A serial dilution of Gli36-derived siEVs enriched with EGFR was detected with the $^{siEV}$PRA. **(B)** The qualitative images are quantified as statistical distributions.

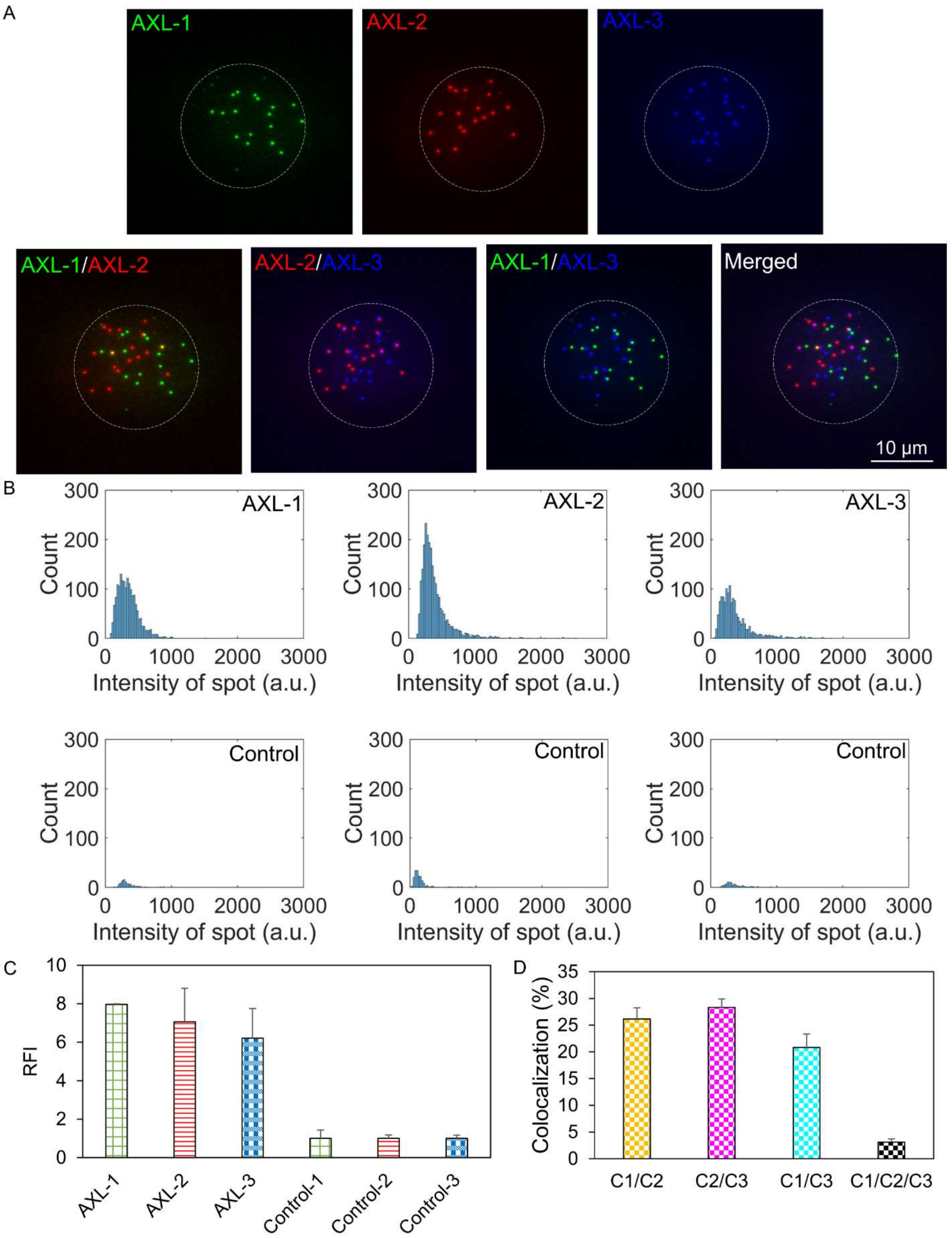

A
AXL-1
AXL-2
AXL-3
AXL-1/AXL-2
AXL-2/AXL-3
AXL-1/AXL-3
Merged
10 μm
B
Count
Intensity of spot (a.u.)
AXL-1
AXL-2
AXL-3
Control
Control
Control
C
RFI
AXL-1
AXL-2
AXL-3
Control-1
Control-2
Control-3
D
Colocalization (%)
C1/C2
C2/C3
C1/C3
C1/C2/C3

**Supplementary Figure 7: Multiple target detection on AXL mRNA**. **(A)** Three regions of the AXL mRNA were detected and multiplexed with the $^{siEV}$PRA on Gli36-derived siEVs. **(B)** The qualitative images for the single region targets are quantified as statistical distributions. **(C)** The qualitative images for the single region targets are quantified as RFI (n = 3, error bars indicate the standard deviation). **(D)** The qualitative images for all three targets are quantified as colocalization efficiencies for the AXL region combinations. C1, C2, and C3 represent the detected biomarkers from left to right (n = 3, error bars indicate the standard deviation).

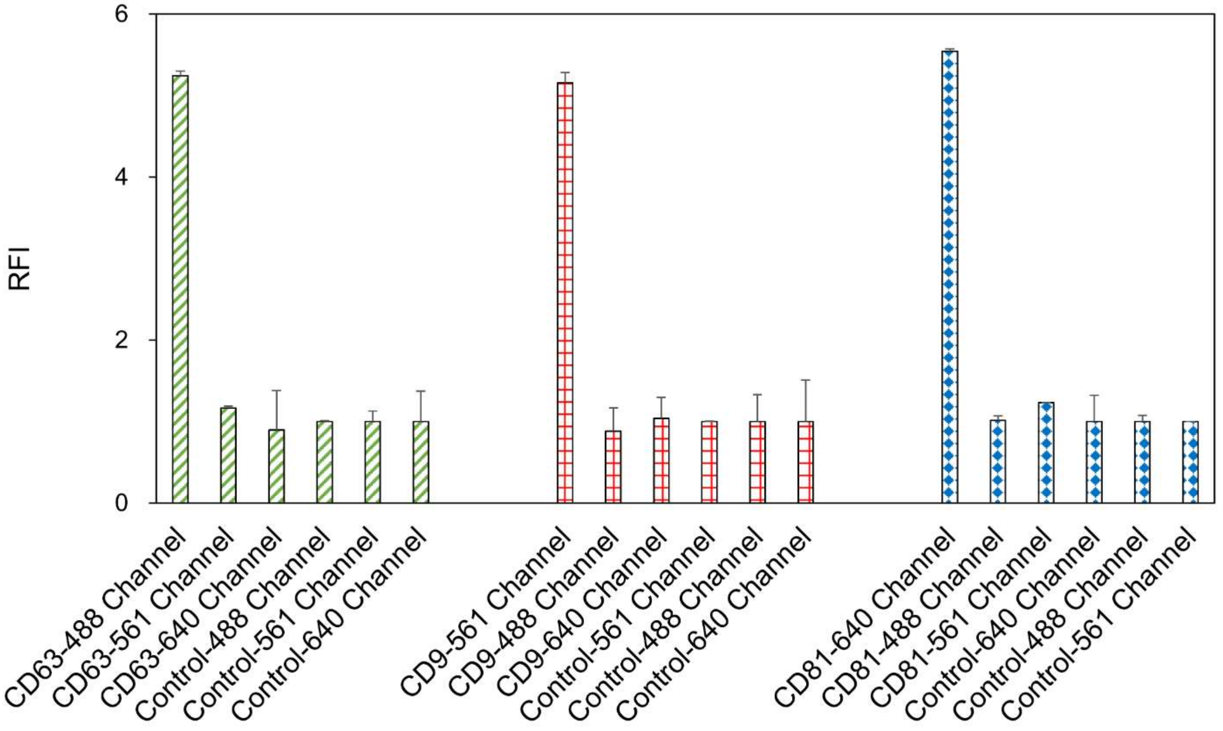


**Supplementary Figure 8: Cross-talk specificity of fluorescent channels.** Fluorescently labeled antibodies, including CD63-488, CD81-640, and CD9-561 were illuminated by different wavelengths and were excited only by their corresponding channel (n = 3, error bars indicate the standard deviation).

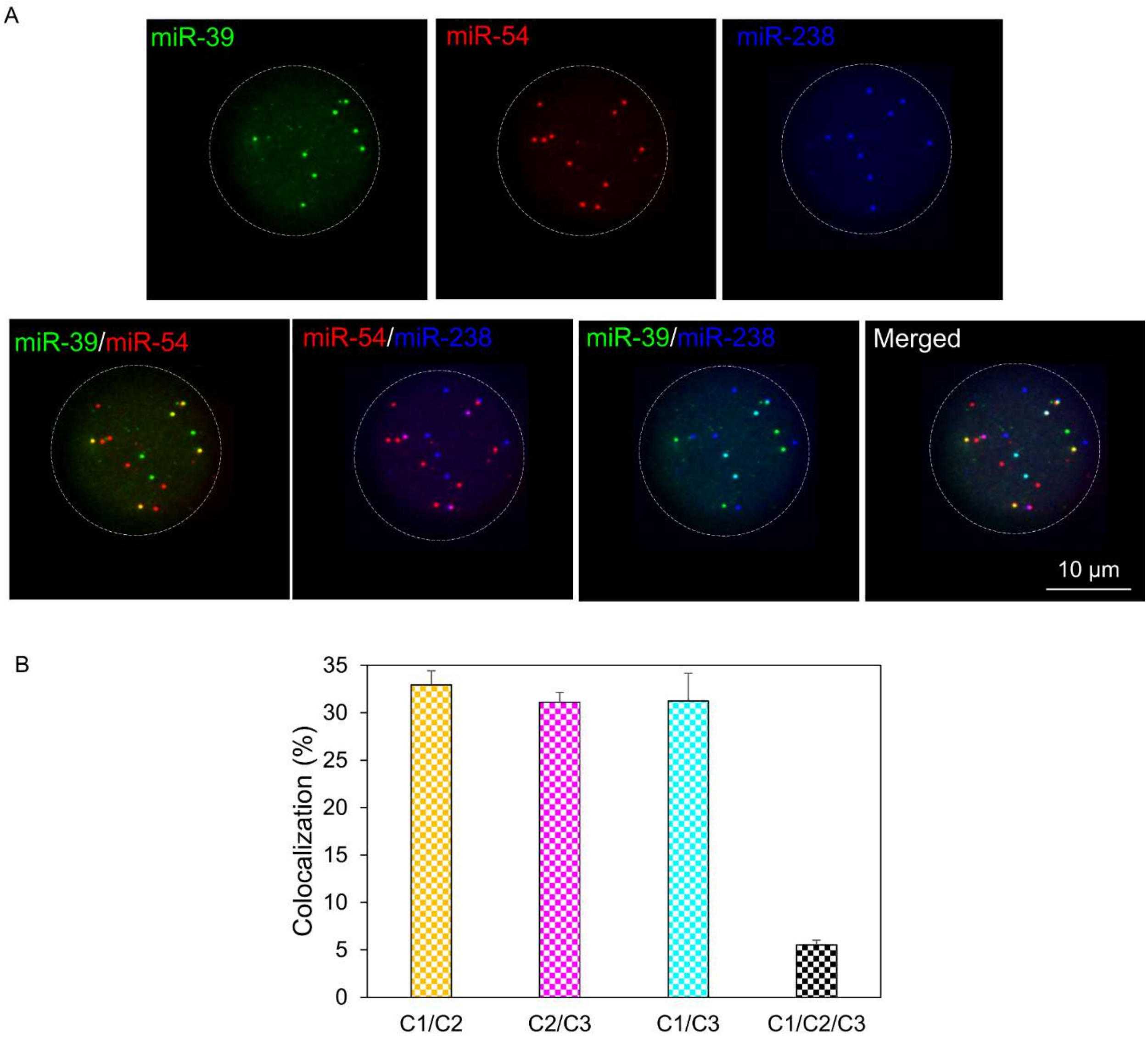


**Supplementary Figure 9: Multiplexed RNA detection. (A)** Multiple miRNA, including miR-39 (green), miR-54 (red), and miR-238 (blue) are multiplexed on siEVs with the $^{siEV}$PRA. **(B)** The qualitative images were quantified as colocalization efficiencies for the miRNA in the Gli36-derived siEVs. C1, C2, and C3 represent the detected biomarkers from left to right (n = 3, error bars indicate the standard deviation).

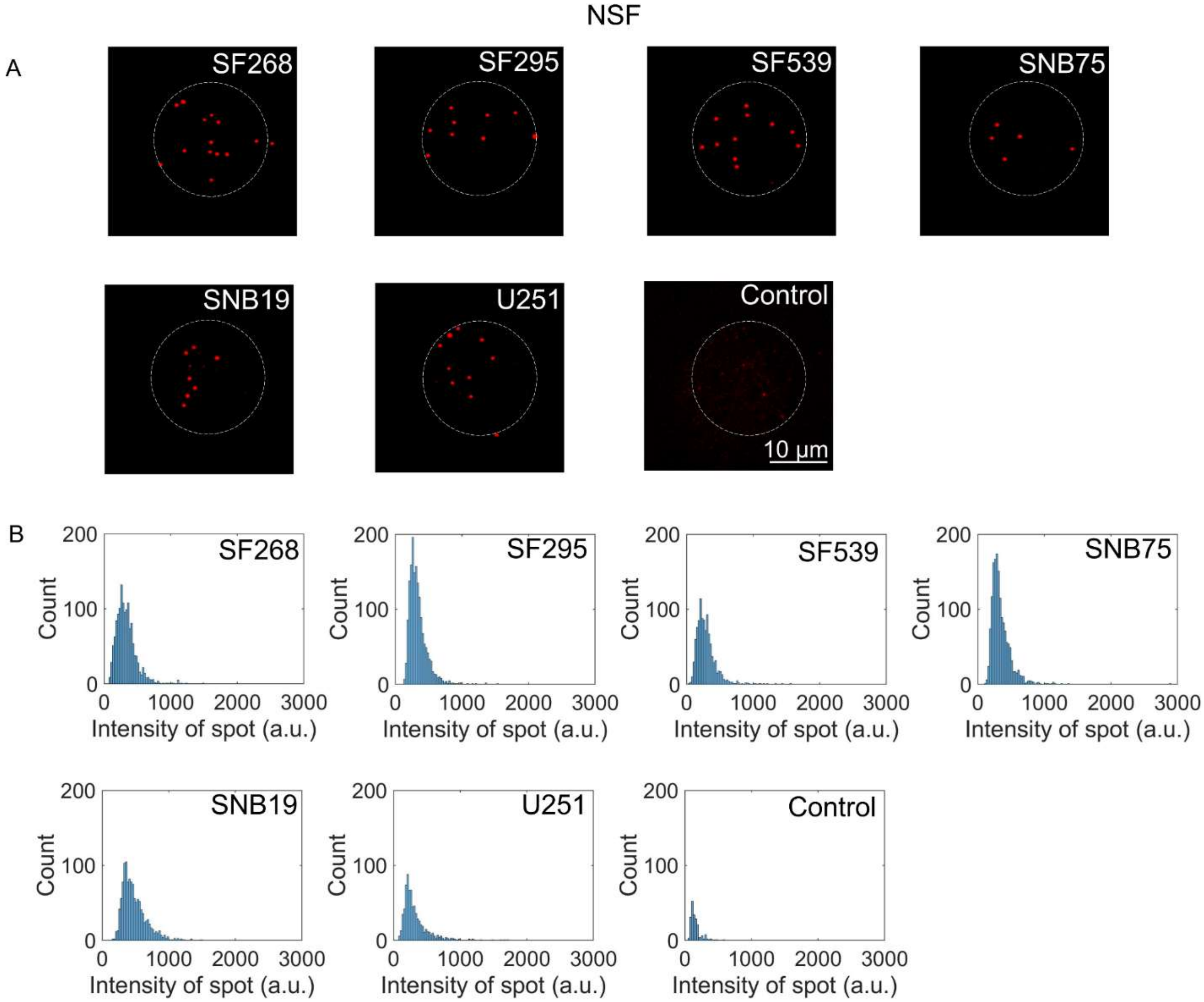


**Supplementary Figure 10: NSF detection with the siEVPRA.** Qualitative images of the siEV detection and statistical distributions for the expression of NSF on siEVs across the six GBM cell lines as detected by the siEVPRA.

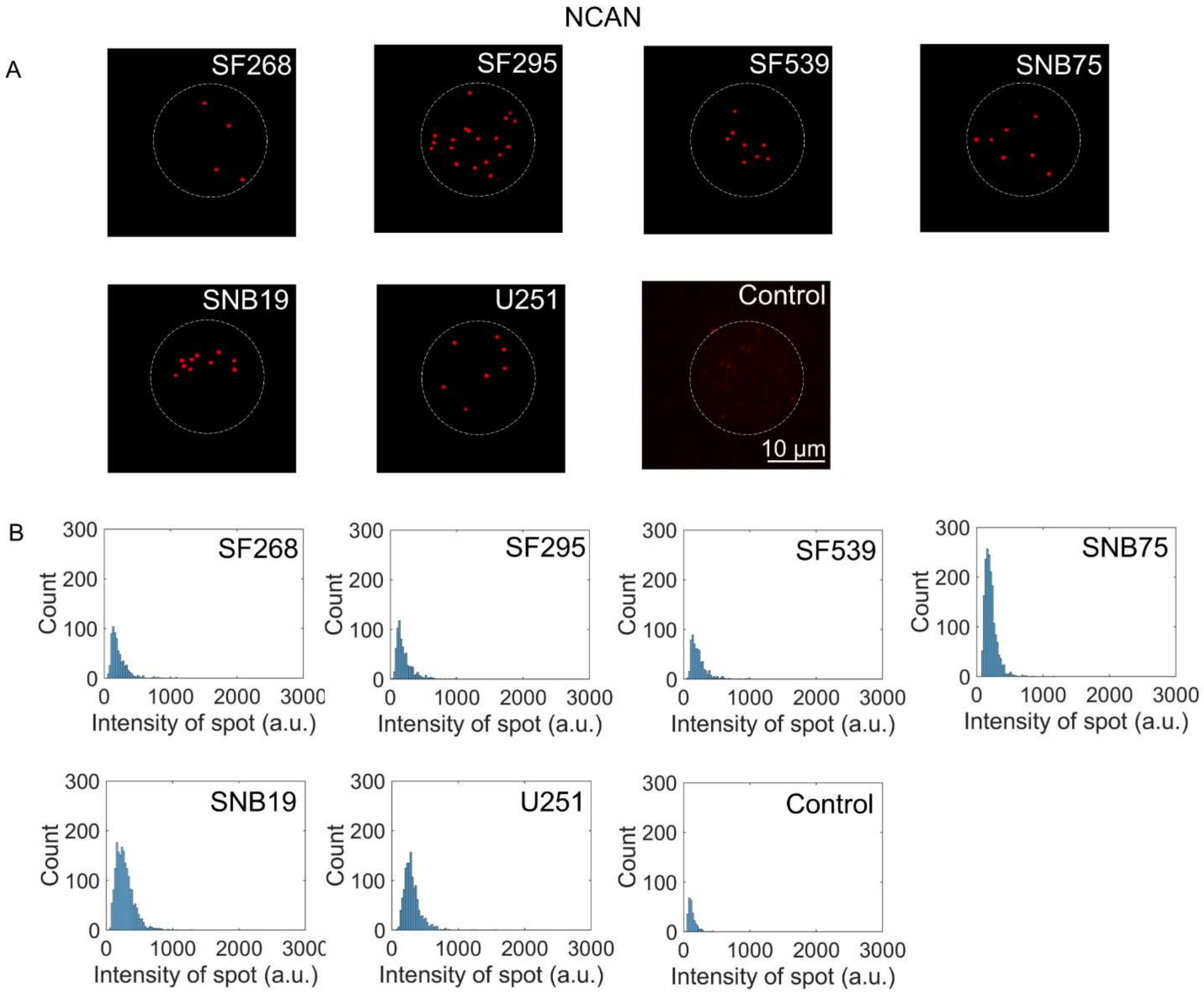


**Supplementary Figure 11: NCAN detection with the $^{siEV}$PRA.** Qualitative images of the siEV detection and statistical distributions for the expression of NCAN on siEVs across the six GBM cell lines as detected by the $^{siEV}$PRA.

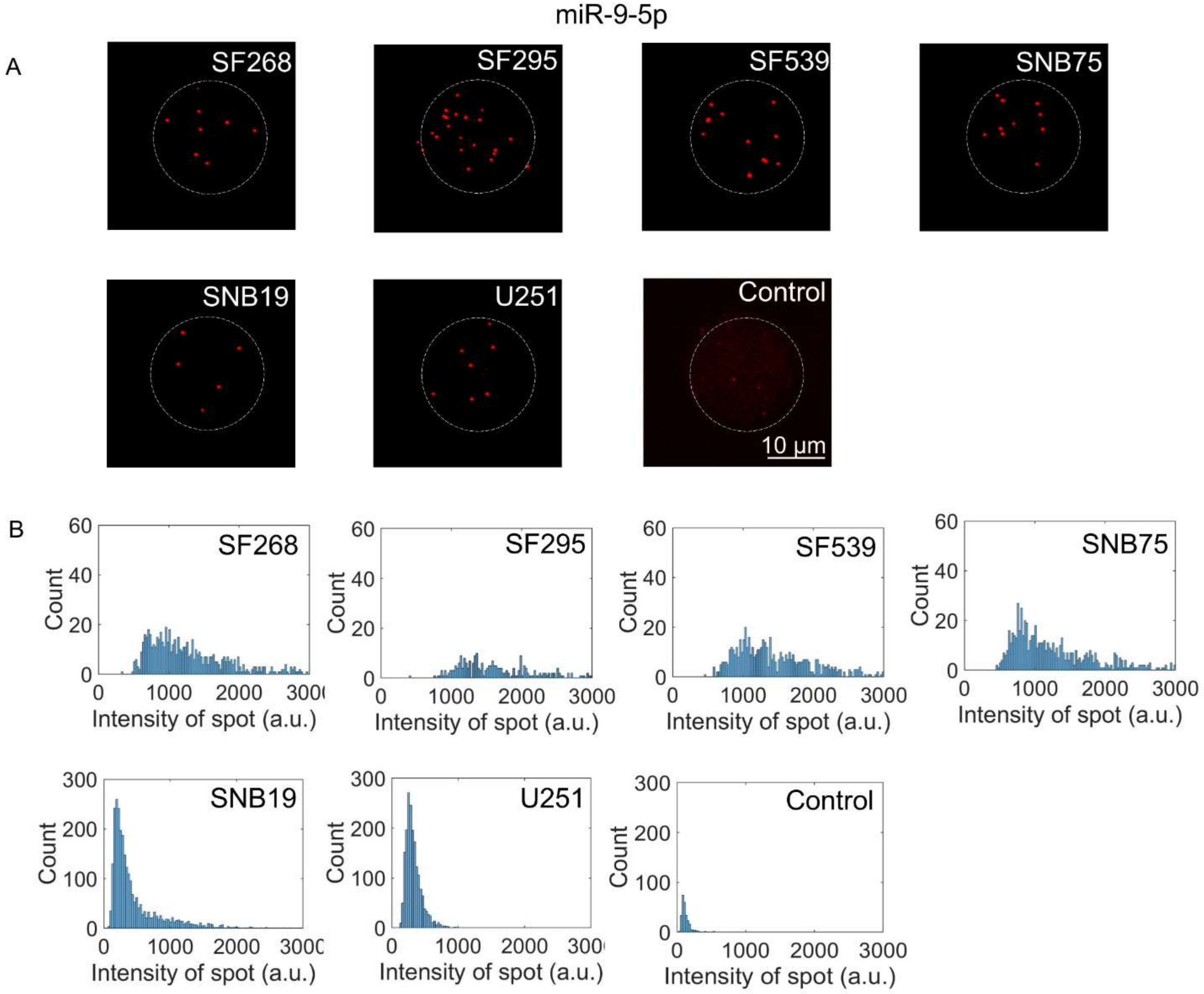


**Supplementary Figure 12: miR-9-5p detection with the $^{siEV}$PRA.** Qualitative images of the siEV detection and statistical distributions for the expression of miR-9-5p on siEVs across the six GBM cell lines as detected by the $^{siEV}$PRA.

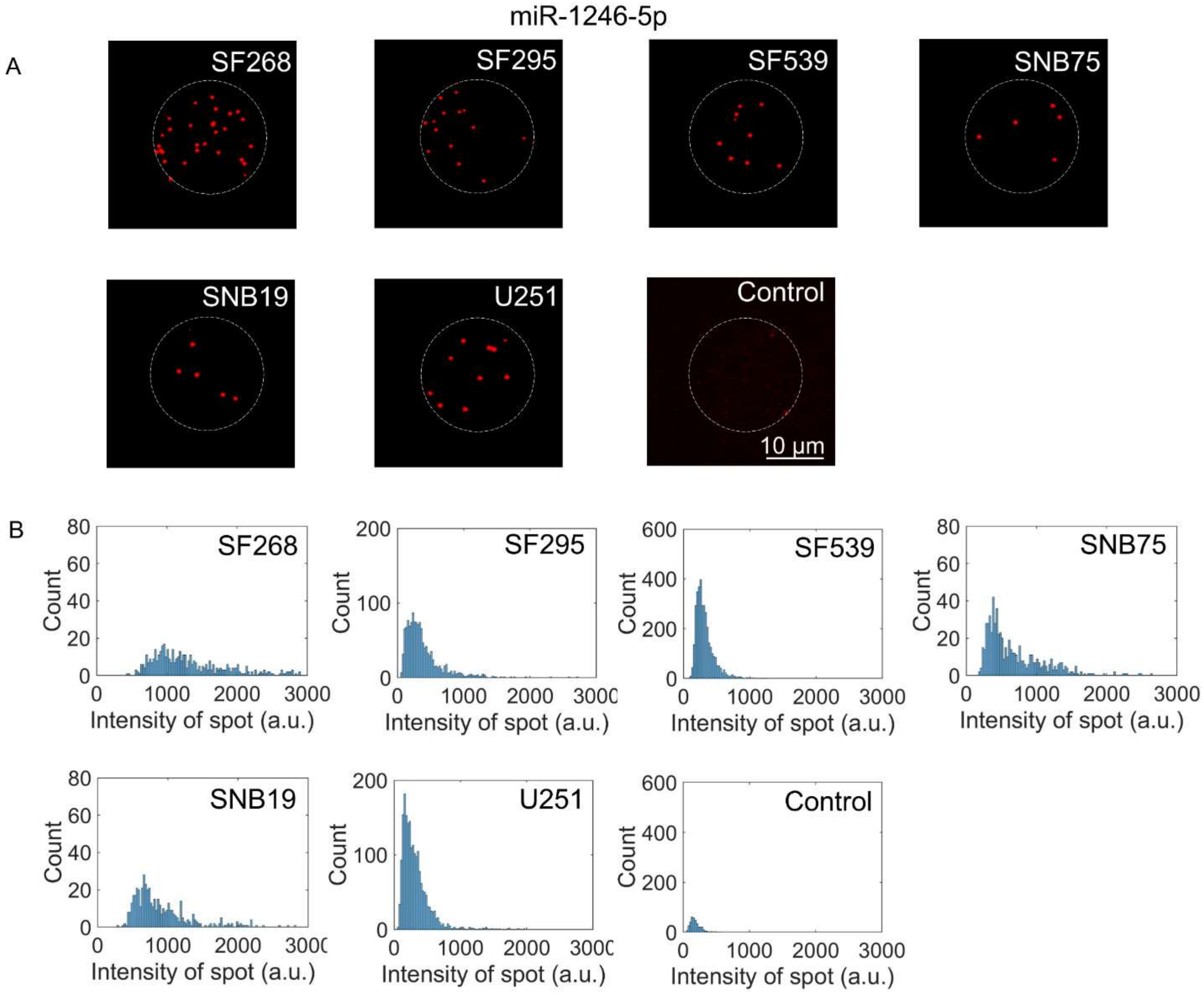


**Supplementary Figure 13: miR-1246-5p detection with the $^{siEV}$PRA.** Qualitative images of the siEV detection and statistical distributions for the expression of miR-1246-5p on siEVs across the six GBM cell lines as detected by the $^{siEV}$PRA.

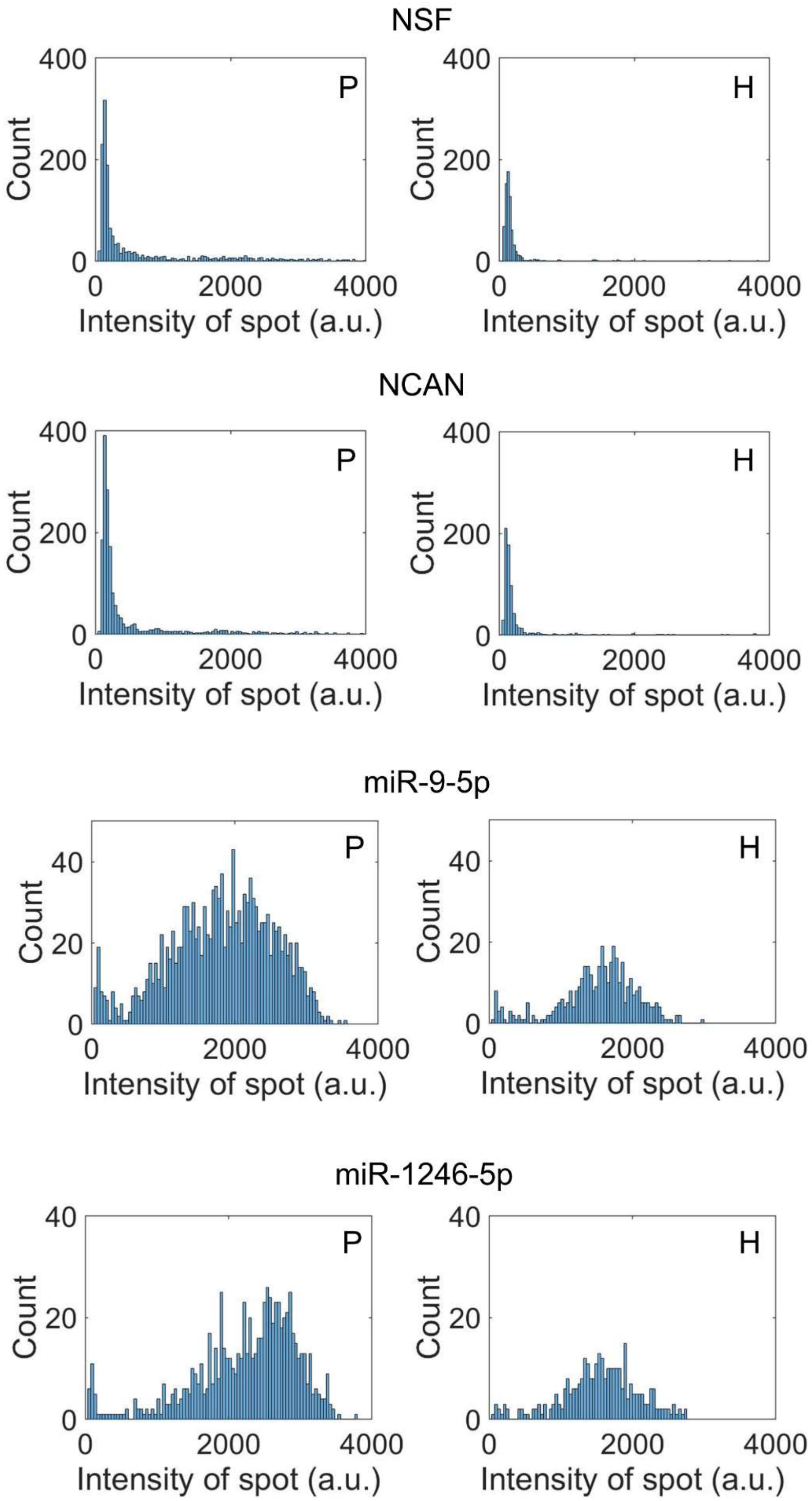
NSF
P
H
NCAN
P
H
miR-9-5p
P
H
miR-1246-5p
P
H
Count
Intensity of spot (a.u.)
0
200
400
20
40
2000
4000

**Supplementary Figure 14: GBM serum-derived EV detection with the $^{siEV}$PRA.** Statistical distributions for the siEV detection of NSF, miR-9-5p, miR-1246-5p, and NCAN for patient (P) and healthy (H) donor samples.
